# Exploring neural correlates of behavioral and academic resilience among children in poverty

**DOI:** 10.1101/2021.09.16.460710

**Authors:** M.E. Ellwood-Lowe, C.N. Irving, S.A. Bunge

**Author notes:** **Correspondence:** Monica E. Ellwood-Lowe, 2121 Berkeley Way, Berkley, CA 94704.

## Abstract

Children in poverty must contend with systems that do not meet their needs. We explored what, at a neural level, helps explain children’s resilience in these contexts. Lower coupling between lateral frontoparietal network (LFPN) and default mode network (DMN)—linked, respectively, to externally- and internally-directed thought—has previously been associated with better cognitive performance. However, we recently found the opposite pattern for children in poverty. Here, we probed ecologically-valid assessments of performance. In a pre-registered study, we investigated trajectories of network coupling over ages 9-13 and their relation to school grades and attention problems. We analyzed longitudinal data from ABCD Study (*N*=8366 children at baseline; 1303 below poverty). The link between cognitive performance and grades was weaker for children in poverty, highlighting the importance of ecologically-valid measures. As predicted, higher LFPN-DMN connectivity was linked to worse grades and attentional problems for children living above poverty, while children below poverty showed opposite tendencies. This interaction between LFPN-DMN connectivity and poverty related to children’s grades two years later; however, it was attenuated when controlling for baseline grades and was not related to attention longitudinally. Together, these findings suggest network connectivity is differentially related to performance in real-world settings for children above and below poverty.

## Introduction

Resources are not equally distributed across a nation’s population; in the United States, the inequity is particularly stark (Zucman, 2019). There is a large body of research focused on the detriments of growing up without as many economic and educational resources (low socioeconomic status, SES). By comparison, far less research has examined how some children in lower-resource contexts are able to adapt and ultimately thrive educationally, exhibiting resilience in the face of structural barriers to success. Measuring children’s brain function is one way to investigate pathways to resilience. For example, one can ask whether children growing up with fewer resources rely on the same neural pathways as their well-off peers to perform well in school, or whether they achieve the same results through alternate means, having adapted meaningfully in order to overcome societal barriers to success. This question has important implications for interventions meant to promote educational success. Should the end-goal of interventions be to make lower-income children’s brain development more closely resemble that of their higher-income peers? If children from different backgrounds achieve success through different neural pathways, a one-size-fits-all approach may not be effective. While single studies alone cannot answer these questions, they can open the door to such lines of inquiry.

A number of brain imaging studies have shown environment-dependent differences in neural recruitment during performance of cognitive tasks (Merz et al., 2019). These studies suggest that the homes, neighborhoods, and schools that form our lived experiences shape our mental and neural processes. This should hardly be surprising, given decades of animal research on experience-dependent brain plasticity (DeFelipe, 2006; Diamond et al., 1964).

In this study, we focus on patterns of brain activation that support behavioral performance in childhood, and where they diverge as a function of family income, a proxy for resource access. One relevant study found that children from higher- and lower-income homes relied on different brain regions to perform well on a working memory task (Finn et al., 2017). Children from higher-income families showed more overall brain activation during task performance: the more they recruited temporal and frontal brain regions, the better they did. Children from lower-income families, on the other hand, showed less activation and did better the *less* they recruited temporal and frontal brain regions. Contrastive findings such as these abound; researchers have typically found differences in frontal and parietal lobe activation as a function of family income, and differences in the ways brain function and structure relates to children’s performance on tasks such as working memory, rule learning, reasoning, and attention (Leonard et al., 2019; Merz et al., 2019; Sheridan et al., 2012).

Another way to test for experience-dependent differences in brain function is with resting state functional MRI (rs-fMRI). This method may more effectively capture the cumulative history of individuals’ experiences and thought patterns (Mackey et al., 2013; Power, Schlaggar, et al., 2014; for a review see Guerra-Carrillo et al., 2014). With rs-fMRI, we measure children’s unconstrained brain activity while they lie in the MRI scanner. The strength of temporal coupling, or so-called functional connectivity, between brain regions—that is, how often they fluctuate in tandem at “rest”—is thought to reflect recent history of coactivation of those regions. Advantages of this method are that it is not influenced by differences in children’s strategy or effort on a particular MRI task, which can be confounds in group comparisons. Functional connectivity measured with rs-fMRI is sensitive to current mental states (e.g., Liston et al., 2009), but also captures brain network connectivity on a broader timescale than a single task performed on a single day.

Here, we focus specifically on children’s resting-state functional connectivity between several brain networks relevant to cognitive and self-referential processing. The lateral frontoparietal network (LFPN) is consistently activated in higher-level cognitive tasks, such as those taxing executive functions or reasoning (Vincent et al., 2008). In contrast, the default mode network (DMN) is more active during internally oriented processing, such as reflecting on one’s self (Raichle et al., 2001), as well as during tasks that require thinking outside of the here-and-now, such as thinking about the past or future (Spreng, 2012). The cingulo-opercular network (CON), sometimes referred to as the “salience network,” has been theorized to serve as an interface between the DMN and LFPN, alerting LFPN to challenges that may require a controlled response, and thus play an important role in switching from the so-called “default” mode to the top-down control mode (Sridharan et al., 2008; Uddin et al., 2011).

The interactions between resting-state networks change as children’s brains develop, though the changes that have been reported depend on the networks and the developmental period in question (Baum et al., 2017; Grayson & Fair, 2017; Marek et al., 2015; Pines et al., 2021). Networks can either become more integrated or more segregated, showing either increased or decreased between-network functional connectivity; network segregation is thought to support network specialization (Baum et al., 2017; Grayson & Fair, 2017; Marek et al., 2015; Pines et al., 2021).

A body of evidence suggests that it is adaptive for LFPN and DMN to be more segregated. First, task-related fMRI studies have shown that stronger activation of the LFPN and stronger deactivation of the DMN is associated with better performance on tasks that require focus on externally presented stimuli (Weissman et al., 2006). Secondly, rs-fMRI research in both adults and children has consistently found that relatively lower functional connectivity between LFPN and DMN is related to better cognitive, emotional, and behavioral outcomes (Chai et al., 2014; DeSerisy et al., 2021; Lopez et al., 2020; Sherman et al., 2014; Whitfield-Gabrieli et al., 2020).

However, LFPN-DMN segregation may not always be adaptive. Mounting evidence suggests that engagement of both the LFPN and DMN is actually beneficial when performing certain kinds of tasks, especially those on which intentional mind-wandering is helpful, such as mentalizing or creative thinking (Christoff et al., 2009; Dixon et al., 2014; Kucyi et al., 2021). Indeed, engaging in deliberate mind-wandering— argued to be distinct from uncontrolled mind-wandering—is thought to help fuel creative insights (Agnoli et al., 2018) and be associated with less reactive emotional processing (Seli et al., 2015). Coactivation of LFPN and DMN has been implicated in performance on a variety of creative thinking tasks (Beaty et al., 2016, 2017; Jaarsveld & Lachmann, 2017), as well as tasks that benefit from drawing on prior knowledge (Spreng & Turner, 2019).

Thus, it is important to understand whether lower LFPN-DMN connectivity is adaptive for all adolescents, whether this relation changes across development, and whether it is adaptive for more ecologically valid measures of cognitive or behavioral performance. In a prior study (Ellwood-Lowe et al., 2021), we found that lower LFPN-DMN connectivity at age 9-11 was linked to better performance on tests of executive functioning and reasoning, but only for children with higher family incomes. To the contrary, children living below poverty—particularly those with specific environmental challenges—tended to show a positive relation between LFPN-DMN connectivity and test performance. This study raises the possibility that children in poverty who more frequently coactivate LFPN-DMN in their day-to-day lives are better poised for academic success, perhaps as a result of engaging in some of the cognitive processes described above. Importantly, however, this study was cross-sectional and only tested performance on laboratory-based cognitive tasks, leaving open questions about the duration and scope of this dissociation between brain development and cognitive performance.

Unlike laboratory-based cognitive tests, children’s grades in school represent their performance in real-world settings, and carry meaningful weight for their future opportunities. On top of this, schools as institutions may be designed in ways that discriminate against children in poverty, particularly those of color (e.g., Darling-Hammond, 2001; Reardon & Owens, 2014; Shedd, 2015). Thus, it is important to understand not only children’s individual test performance, but how this relates to their performance in real institutional settings, which often pose additional or alternative barriers to success. While cognitive test performance and grades are typically correlated (e.g., Best et al., 2011; Cowan, 2014; St Clair-Thompson & Gathercole, 2006; Willoughby et al., 2019), we hypothesized that they may be less so for children in poverty. With regard to brain function, it is possible that the dissociation between children above and below poverty for LFPN-DMN connectivity would not be observed for this more real-world measure of performance in institutional contexts. However, given prior evidence that this dissociation may have something to do with adapting in the face of structural barriers (Ellwood-Lowe et al., 2021), we predicted that it would also be present for children’s grades in school.

Another more ecologically valid assessment of performance in the real-world is that of children’s attention problems. Attention problems can pervade many aspects of children’s lives, from their performance in school to self-esteem to relationships with peers, teachers, and family members (Harpin, 2005). Importantly, less LFPN-DMN connectivity has been linked to fewer attention problems (Whitfield-Gabrieli et al., 2020). Thus, one possibility is that children in poverty with higher LFPN-DMN connectivity are similarly at elevated risk of other problems, such as attentional difficulties. Alternatively, the dissociation found previously between LFPN-DMN connectivity and cognitive test performance may also be observed for attentional problems.

Given the purported role of the CON in toggling between internally and externally focused attention, the interplay between CON and the DMN and LFPN may also be related to grades and/or attentional problems, in a way that may depend on children’s environments. Previous research has shown that CON network dynamics are related to performance on IQ tests (Hilger et al., 2017). Moreover, in a previous exploratory analysis, we observed a trend such that higher CON-DMN connectivity was directionally related to better test scores for children in households above poverty and worse scores for children in households below poverty. Thus, it is of interest to further explore patterns of CON connectivity with the DMN and LFPN.

### Present study

We sought to test three primary aims related to LFPN-DMN connectivity, grades, and attention, as well as two secondary aims related to CON connectivity (see Supplementary Table 1). Data came from a baseline assessment when children were approximately ten years old (T0) and a follow-up assessment two years later (T2). By using these measures, we sought to better understand the neural basis of children’s resilience with regard to societal constraints. Importantly, these measures assess functioning over a broader timeframe than their score on a set of tests completed on a single day. We delineate our aims and pre-registered predictions below, along with relevant exploratory follow-up analyses.

#### Primary Aim 1

First, we sought to characterize longitudinal changes in LFPN-DMN resting-state functional connectivity, and test whether they varied as a function of poverty status. Based on prior findings (Baum et al., 2017; Sherman et al., 2014), we predicted that the LFPN and DMN would generally become less coupled over early adolescence for children above poverty. Given that we had not previously observed baseline differences between the groups in LFPN-DMN connectivity (Ellwood-Lowe et al., 2021), we hypothesized that children below poverty would show a similar decline in connectivity to these higher-income peers (H1).

##### Exploratory

Because our previous study showed that lower-performing children below poverty showed lower LFPN-DMN connectivity than other children, we conducted follow-up analyses investigating whether trajectories of network connectivity differ as a function of children’s cognitive test scores and their poverty status. While not pre-registered, this exploratory analysis tested our pre-registered aim to consider three equally plausible hypothetical patterns of data: convergence, further divergence, or stability of differences in functional connectivity between lower-performing children below poverty and other children.

#### Primary Aim 2

Second, to characterize concurrent and longitudinal relations between our behavioral variables, we sought to assess the extent to which laboratory-based cognitive tests are linked to more ecologically valid measures of performance for children above and below poverty. In particular, we examined cross-sectional and longitudinal associations between test scores and children’s grades in school, for children above and below poverty.

##### Concurrent

We predicted that children’s performance on cognitive tests at baseline would be associated with their grades in school (H2a). However, we hypothesized that the concurrent association between cognitive performance and grades may be weaker for children in poverty, who face many potential barriers to academic success (H2b). If this were the case, we expected to find an interaction for the relation between concurrent cognitive performance and grades as a function of poverty status.

##### Longitudinal

Given Cattell’s classic hypothesis that current cognitive functioning supports the acquisition of crystallized knowledge, we planned to test whether cognitive test performance at T0 predicted grades at T2, controlling for grades at T0. We hypothesized that the longitudinal association between cognitive performance and grades may be weaker for children in poverty (H3). If this were the case, we expected to find an interaction for the relation between cognitive performance and grades as a function of poverty status.

#### Primary Aim 3

Third, we sought to assess the extent to which LFPN-DMN connectivity is linked to more ecologically valid measures of cognitive performance for children above and below poverty. Specifically, we planned to examine cross-sectional and longitudinal associations between LFPN-DMN connectivity and grades and attention problems, for children above and below poverty.

##### Grades, concurrent

We hypothesized that LFPN-DMN connectivity would be differentially associated with academic performance for children above and below poverty. Our primary prediction was a concurrent relation (H4): higher LFPN-DMN connectivity would be associated with higher grades for children in poverty and lower grades for children above poverty, mirroring our findings for test scores.

##### Grades, longitudinal

Based on prior research showing longitudinal but not concurrent brain-behavior relations, we also sought to test the hypothesis that LFPN-DMN connectivity supports knowledge acquisition (H5). We predicted that LFPN-DMN connectivity would be longitudinally associated with grades, controlling for grades at T0.

##### Attention, concurrent (Exploratory)

As described next, we predicted a longitudinal relation involving this measure; by contrast, we did not have a strong predictions for a cross-sectional relation. However, for the sake of completeness, we also ran an exploratory cross-sectional analysis testing concurrent relationships between LFPN-DMN connectivity and attention problems at T0 for children above and below poverty.

##### Attention, longitudinal

Given prior evidence that stronger LFPN-DMN connectivity was linked to more attention problems longitudinally but not concurrently (Whitfield-Gabrieli et al., 2020), we sought to assess whether stronger LFPN-DMN connectivity is associated with greater attention problems longitudinally for children above poverty, controlling for attention problems at T0. If this is a general phenomenon, we would expect children below poverty to show the same relation (H6a). However, given our prior findings regarding differential brain-behavior relations for cognitive performance, children below poverty could conceivably show the opposite pattern (H6b).

#### Secondary Aims

Our secondary aims focused on CON-DMN and CON-LFPN given that CON is theorized to serve as an intermediary the DMN and LFPN, enabling switching attention between internally and externally guided mental states. Overall, we predicted that both networks would decrease in coupling across early adolescence, though potentially less so for children in poverty. Further, we predicted lower CON-LFPN would be related to children’s grades in school. Finally, we predicted that lower CON-DMN and CON-LFPN would be related to fewer attention problems, though this effect might be stronger for children in poverty. (See Supplement for full description of aims and associated analyses.)

## Methods

### Parent study

Data were drawn from the Adolescent Brain Cognitive Development (ABCD) study, which was designed to recruit a large cohort of children who closely represented the United States populations (http://abcdstudy.org; see Garavan et al., 2018). The ABCD study is a multisite, longitudinal study intended to run for at least 10 years following 11,878 children, recruited at ages 9-11, into late adolescence. A wide variety of data are collected on each youth including mental and physical health assessments, behavioral data, imaging data, and more. This study was approved by the Institutional Review Board at each study site, with centralized IRB approval from the University of California, San Diego. Informed consent and assent were obtained from all parents and children, respectively. We include data from two timepoints: T0 (baseline assessment; ages 8.9-11.1) and T2 (two-year follow-up; ages 10.6-13.6). We note that while there is a range of ages represented at each timepoint, overall ages were relatively tight; most children were almost precisely ten years old at T0 and twelve years old at T2 (see Table 1). Children also completed behavioral measures at an interim one-year follow-up (T1); while not the focus of the current study, we report additional analyses with T1 data in the Supplement.

**Table 1.**
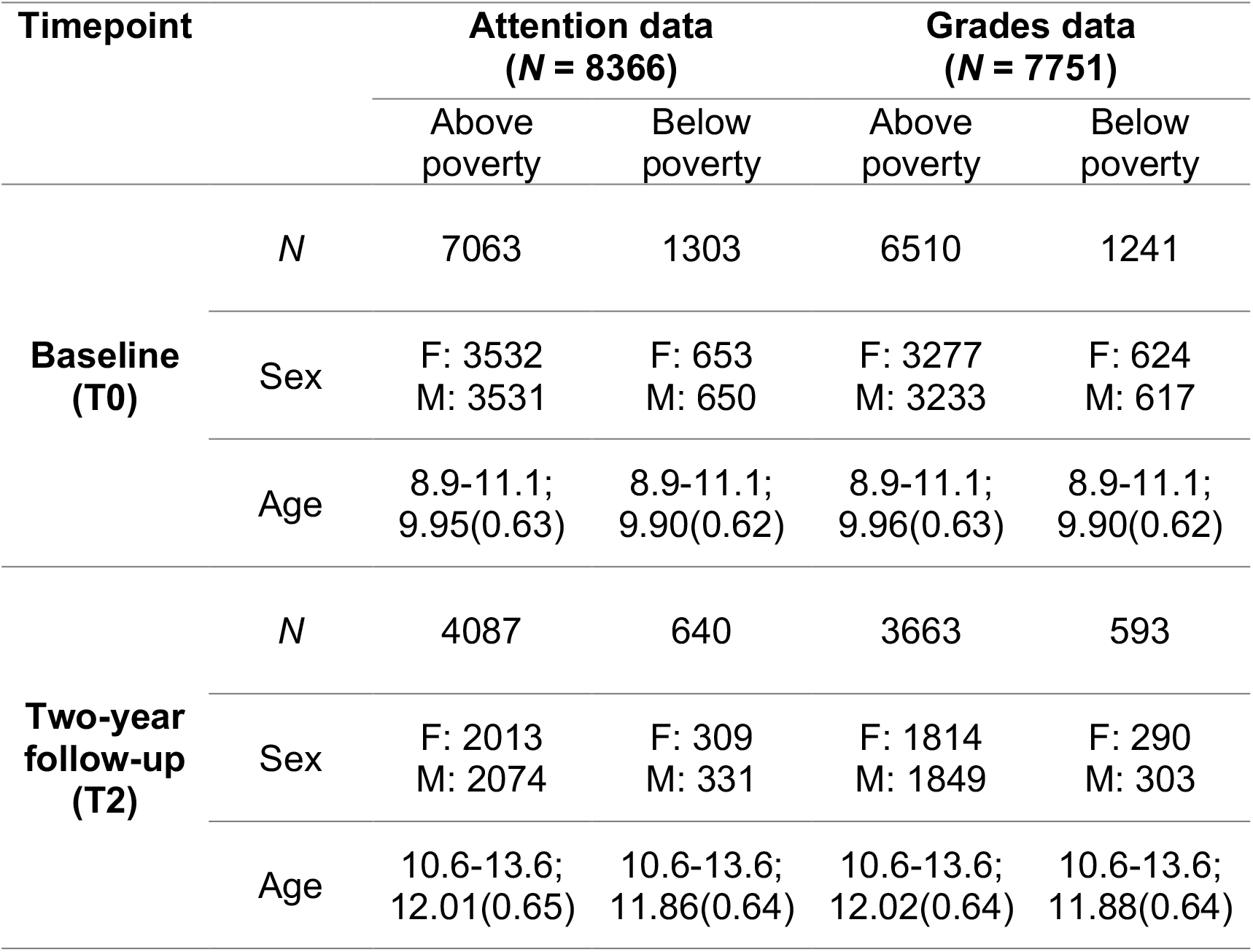
Sample sizes, age (range; mean(SD)), and parent-reported child sex for children above and below poverty, at T0 and T2. Sample sizes differ slightly for those analyses focusing on grades and those focusing on attention, based on the number of children providing usable data.

### Present study

Planned analyses were pre-registered prior to data access (https://aspredicted.org/QWQ_C5N; https://aspredicted.org/NTG_RRB) and analysis scripts are available on the Open Science Framework (https://osf.io/gcjn8/?view_only=d0f098d6a8ab47d5bf0bbb290141bbd3). The original data are available with permissions on the NIMH Data Archive (https://nda.nih.gov/abcd). All deviations from the initial analysis plan are fully described in the Supplement.

Children from the full sample were excluded if they did not provide data on any of the measures used in our analyses. Specifically, children were excluded from analyses if their caregiver did not provide information about family income at baseline (*N* = 1,146), if they were missing rs-MRI data at baseline (*N* = 101) or their rs-fMRI data did not meet ABCD’s usability criteria (*N* = 2,390), if they did not provide usable cognitive test score data at baseline (*N* = 237), or if there was no information about the child’s age, sex, or family ID (used to track whether participants were siblings; *N* = 2; see Supplement for exclusion flow-charts for T0 and T2). After these initial exclusions, children who had data related to attention problems were included in analyses relevant to attention (*N* = 8366), and those with data related to grades in school were included in analyses relevant to grades (*N* = 7751; see Table 1). Because the former group included slightly more children, we used this sample when conducting analyses that did not include behavioral data.

On average, children who were excluded were more likely to be younger, be male, have more attention problems, have lower grades, and have lower cognitive test scores. While children in poverty were also more likely to be excluded, these differences between included and excluded participants were observed for both children above and below poverty (see Supplement).

We estimated poverty status at baseline for each child based on their combined family income bracket, the number of people living in the home, and the average supplemental poverty level for the study sites included in the sample, as in our previous work (Ellwood-Lowe et al., 2021). Based on the factors used to estimate poverty status, we considered children to be living below the poverty line if they were living in a household of 4 with a total income of less than $25,000, or a household of 5 or more with a total income of less than $35,000 at T0 (Table 1). We did not consider whether poverty status changed after T0.

### Behavioral measures

Children’s cognitive performance was measured at T0 with a cognitive test battery that included measures from the NIH Toolbox (http://www.nihtoolbox.org). The NIH Toolbox Fluid Cognition composite measure includes two tests of working memory (Picture Sequence Memory Test, List Sorting Working Memory Test), two tests of executive functions that tap into cognitive flexibility and inhibitory control (Dimensional Card Sort and Flanker tasks), and one test of processing speed (Pattern Comparison Processing Speed Test). The administered test battery also included the Matrix Reasoning Task from the Wechsler Intelligence Test for Children-V (WISC-V), a measure of abstract reasoning (Wechsler, 2014). More details on each of these tests and their administration in the current study is described elsewhere (Luciana et al., 2018).

Attention and behavioral problems were measured with the Attention subscale of the Child Behavior Checklist (CBCL). The CBCL is a standardized form which is used to characterize children’s externalizing and internalizing behaviors (Achenbach & Ruffle, 2000). From the initial baseline assessment onwards, parents completed an automated version of the CBCL annually, reporting on their child’s behavior over the past six months (Barch et al., 2018). Each item on the CBCL was rated using a three-point rating scale: “not true,” “somewhat or sometimes true,” “very true or often true.” There were 11 items in the attention subscale. We used the mean of all items at T0 and T2, separately, for each child. Higher scores indicated more attentional and behavioral problems.

Children’s academic performance was measured via parent-reported grades in the ABCD Longitudinal Parent Diagnostic Interview for DSM-5 Background Items Full (KSAD). Parents were asked what kind of grades their child received on average: 1= As/excellent, 2 = B’s/Good; 3 = C’s/Average; 4 = D’s/Below Average; 5 = F’s/Struggling a lot; 6 = ungraded, -1 = unapplicable. This question was asked at T0 and T2.

### MRI Scan Procedure

Scans were collected on one of three types of 3T scanners (Siemens, Philips, or GE) with an adult-size head coil. Resting state scans were completed at T0 and T2. Scans were typically completed on the same day as the cognitive battery, but could also be completed at a second testing session. After completing motion compliance training in a simulated scanning environment, participants first completed a structural T1-weighted scan. Next, they completed three to four five-minute resting state fMRI scans, during which they were instructed to lay with their eyes open while viewing a crosshair on the screen. The first two resting state scans were completed immediately following the T1-weighted scan; children then completed two other structural scans, followed by one or two more resting state scans, depending on the protocol at each specific study site.

Scan parameters were optimized to be compatible across scanners, allowing for maximal comparability across the 19 study sites. T1-weighted scans were collected axially with 1mm^3^ voxel resolution, 256 × 256 matrix, 8º flip angle, and 2x parallel imaging. Other scan parameters varied by scanner platform (Siemens: 176 slices, 256 × 256 FOV, 2500 ms TR, 2.88 ms TE, 1060 ms TI; Philips: 255 slices, 256 × 240 FOV, 6.31 ms TR, 2.9 ms TE, 1060 ms TI; GE: 208 slices, 256 × 256 FOV, 2500 ms TR, 2 ms TE, 1060 ms TI). fMRI scans were collected axially with 2.4mm^3^ voxel resolution, 60 slices, 90 × 90 matrix, 216 × 216 FOV, 800ms TR, 30 ms TE, 52º flip angle, and 6 factor MultiBand Acceleration. Head motion was monitored during scan acquisition using real-time procedures (fMRI Integrated Real-time Motion Monitor; Dosenbach et al., 2017) to adjust scanning procedures and collect additional data as necessary (Casey et al., 2018). This prospective motion correction procedure significantly reduces scan artifacts due to head motion (Hagler et al., 2019), which are known to affect functional connectivity estimates (Power et al., 2015; Satterthwaite et al., 2013).

Structural and functional images underwent automated quality control procedures (including detecting excessive movement and poor signal-to-noise ratios) and visual inspection and rating (for structural scans) of images for artifacts or other irregularities (described in Hagler et al., 2019). Participants were excluded if they did not meet the ABCD study’s quality control criteria for both structural and functional scans.

Specifically, structural scans were excluded if there was evidence of severe inaccuracy of structural surface reconstruction based on any of the five categories: motion, intensity inhomogeneity, white matter underestimation, pial overestimation, or magnetic susceptibility artifact. Functional scans were excluded if they did not meet motion criteria, described further below. In total, 2,390 participants were excluded based on these criteria; in the ABCD dataset, data that passes quality control checks are denoted as IMGINCL_RSFMRI_INCLUDE==1. Excluded participants differed meaningfully from the rest of the sample (see above; more information in Supplement).

Altogether, there was a four-step process for reducing the effect of head motion on rs-fMRI results. First there was real-time head motion monitoring and correction, as described above. Second, there was a thorough and systematic check of scan quality in collaboration with ABCD’s Data Analysis and Informatics Center. Third, signal from motion timecourses was regressed out during preprocessing, and frames with greater than 0.2mm of framewise displacement were excluded from calculations altogether, as were time periods with less than five contiguous low-motion frames. Fourth, a final censoring procedure was employed to identify potential lingering effects of motion by excluding any frames with outliers in spatial variation across the brain (Hagler et al., 2019). In combination, these procedures reduce motion artifacts (Power, Mitra, et al., 2014).

### Resting state fMRI preprocessing

Data preprocessing was carried out using the ABCD pipeline and carried out by the ABCD Data Analysis and Informatics Core; more details are reported by Hagler et al. (2019). The approach for calculating between-network connectivity was decided by the ABCD consortium based on best practices for the field; we use the publicly available processed data in these analyses, but outline the processing steps described in Hagler et al. (2019) here.

Briefly, T1-weighted MR images were corrected for gradient nonlinearity distortion and intensity inhomogeneity, and rigidly registered to a custom reference brain (Friston et al., 1995). These images were run through FreeSurfer’s automated brain segmentation to derive white matter, ventricle, and whole brain ROIs. Resting state fMRI data were first corrected for head motion, displacement estimated from field map scans, B0 distortions, and gradient nonlinearity distortions, and registered to the structural images using mutual information. Initial scan volumes were removed, and each voxel was normalized and de-meaned. Signal from estimated head motion timecourses (including six motion parameters, their derivatives, and their squares), quadratic trends, and mean timecourses of white matter, gray matter, and whole brain, plus first derivatives, were regressed out, and frames with more than 0.2mm displacement were excluded. The data then underwent temporal bandpass filtering (0.009 – 0.08 Hz).

To derive between-network connectivity metrics, data were then projected onto each individual’s cortical surface space, and ROIs were labeled using Freesurfer’s anatomically-defined parcellations (Desikan et al., 2006; Destrieux et al., 2010). ROIs were assigned to one of 13 functionally defined networks, as defined by Gordon et al (2016); for a list of every ROI and its network assignment, see Supplementary Table 5 in Hagler et al. (2019). To calculate between-network connectivity, the timecourse of each ROI in one network was correlated with the timecourse of each ROI in the other network, in a pairwise fashion. These pairwise correlations were z-transformed and averaged to calculate between-network connectivity across all networks. Thus, this between-network connectivity metric represents the average correlation of each ROI in one network with each ROI in the other network. Here, we examined between-network LFPN-DMN, CON-DMN, and CON-LFPN connectivity.

### Analyses

Analyses were performed using the software package R (version 4.0.2; R Core Team, 2017). To determine significance in our models, we performed nested model comparison. In all cases, we compared models without the inclusion of the variable of interest to models with this variable included; we calculated whether the variable of interest contributed significantly to model fit using the *anova* function for likelihood ratio test model comparison. For models in which the dependent variable was continuous, we performed linear mixed effects models using the *lme4* package (Bates et al., 2015); for models in which this variable was an ordered factor (e.g., grades), we performed cumulative link mixed models using the *ordinal* package (Christensen, 2018).

To determine whether to include potential covariates in our model, we tested whether each of the following variables contributed significantly to model fit: (1) a random intercept of study site, (2) a random intercept of families, (3) a fixed effect of sex, (4) a fixed effect of child age, and (5) a fixed effect of head motion (mean framewise displacement) during the resting state scan. All covariates besides age contributed to model fit at a level of *p* < .05 and were thus retained in final models.

#### Longitudinal changes in brain network connectivity

We examined network changes over early adolescence, and whether these changes differed as a function of poverty status. We performed three separate linear mixed effects models testing the interaction of timepoint (T0, T2) with poverty status (above, below), in association with (1) LFPN-DMN connectivity, (2) CON-DMN connectivity, and (3) CON-LFPN connectivity.

#### Behavioral measures

We assessed the concurrent and longitudinal relations between cognitive test performance and grades, and tested whether the relation varied as a function of poverty status. To this end, we conducted cumulative link mixed models to test grades in association with an interaction between poverty status and test performance. We had preregistered an analysis plan using linear mixed effects models to test these relations. However, because grades are a categorical ordered variable, cumulative link mixed models are more appropriate. Thus, we report the latter analyses for all tests including grades as an outcome variable. Results are not meaningfully different when performing the pre-registered linear mixed effects models.

#### Functional connectivity in relation to grades and attention

We investigated the relation between children’s grades and LFPN-DMN connectivity at T0 and T2. We performed two separate cumulative link mixed models to characterize the relation between children’s academic performance and LFPN-DMN connectivity. The first model tested this relation at T0, with an interaction between LFPN-DMN connectivity and poverty status. The second model tested this relation longitudinally, to see whether LFPN-DMN at T0 and its interaction with poverty status related to children’s academic performance at T2, respectively, when controlling for children’s academic performance at T0.

Similarly to our analyses focused on grades, we examined the relation between children’s attention problems and LFPN-DMN connectivity at T0 and T2. While the concurrent association was not pre-registered, this exploratory analysis allowed us to test for the stability of this association over time. As attention was a continuous variable, we performed two separate linear mixed effects models testing the interaction between LFPN-DMN connectivity with poverty status, in association with (1) attention problems at T0 and (2) attention problems at T2, controlling for children’s attention at T0.

In preregistered secondary analyses, we also examined relations between these behavioral measures and CON network connectivity.

#### Exploratory analyses

We conducted several additional follow-up analyses, following a similar analytic approach to those above, for which we did not have strong predictions but that provide a more complete picture of the relations among key variables. These additional analyses are reported in the main text and Supplementary Materials. Those that were not preregistered are indicated as exploratory.

## Results

### Aim 1: Longitudinal changes in network connectivity

#### LFPN-DMN connectivity over adolescence

As a primary analysis, we first, we examined how network connectivity changed between T0 and T2 in LFPN-DMN resting-state functional connectivity. We predicted that these networks would become less coupled between ages 9 and 13. However, there was no significant change in LFPN-DMN connectivity across the group over the course of the two years, *B* = 0.001, SD = 0.001, χ^2^ (2) = 1.01, *p* = .605. We also predicted that children above and below poverty would not exhibit differential trajectories of change; in line with this prediction, there was no evidence for a significant interaction of poverty status with timepoint, interaction: *B* = -0.001, SD = 0.002, χ^2^ (1) = 0.2, *p* = .651. Rather, we observed marked individual variability in the slope and magnitude of change over time (Supplementary Figure 1). An exploratory analysis further suggested that this effect was not meaningfully moderated by children’s cognitive test performance at T0 (see Supplement).

#### CON-DMN and CON-LFPN connectivity over adolescence

As secondary analyses, we tested change in connectivity between our two CON networks of interest. As predicted, both CON-DMN and CON-LFPN both decreased in connectivity between T0 and T2. Also as predicted, the decrease in CON-DMN connectivity was attenuated for children in poverty; change in CON-LFPN connectivity did not meaningfully differ between groups (see Supplement).

### Aim 2. Associations between cognitive test performance and children’s grades in school and attention problems

As primary analyses, we next tested cross-sectional and longitudinal relations between parent-reported grades and cognitive test scores. We predicted that children’s performance on cognitive tests at T0 would be concurrently related with their grades in school. Furthermore, we predicted that this relation would differ as a function of poverty status, with children in poverty showing a weaker relation.

#### Concurrent relations between test scores and grades

As predicted, higher scores on the NIH composite were related to better grades in school concurrently, *B* = - 0.08, SD = 0.004, χ^2^ (2) = 752.25, *p* < .001 (Figure 1). Additionally, as predicted, the relation between cognitive performance and academic achievement differed as a function of poverty status, *B* = 0.03, SD = 0.01, χ^2^ (1) = 24.31, *p* < .001. Follow-up analyses showed that the relation was significant for children above and below poverty (below poverty: *B* = -0.04, SD = 0.01, χ^2^ (1) = 63.70, *p* < .001; above poverty: *B* = -0.08, SD = 0.004, χ^2^ (1) = 688.89, *p* < .001), although the effect was stronger for children above poverty. Thus, children’s performance on cognitive tests at T0 is concurrently associated with their grades in school, albeit less so for children in poverty than those above poverty.

**Figure 1.**
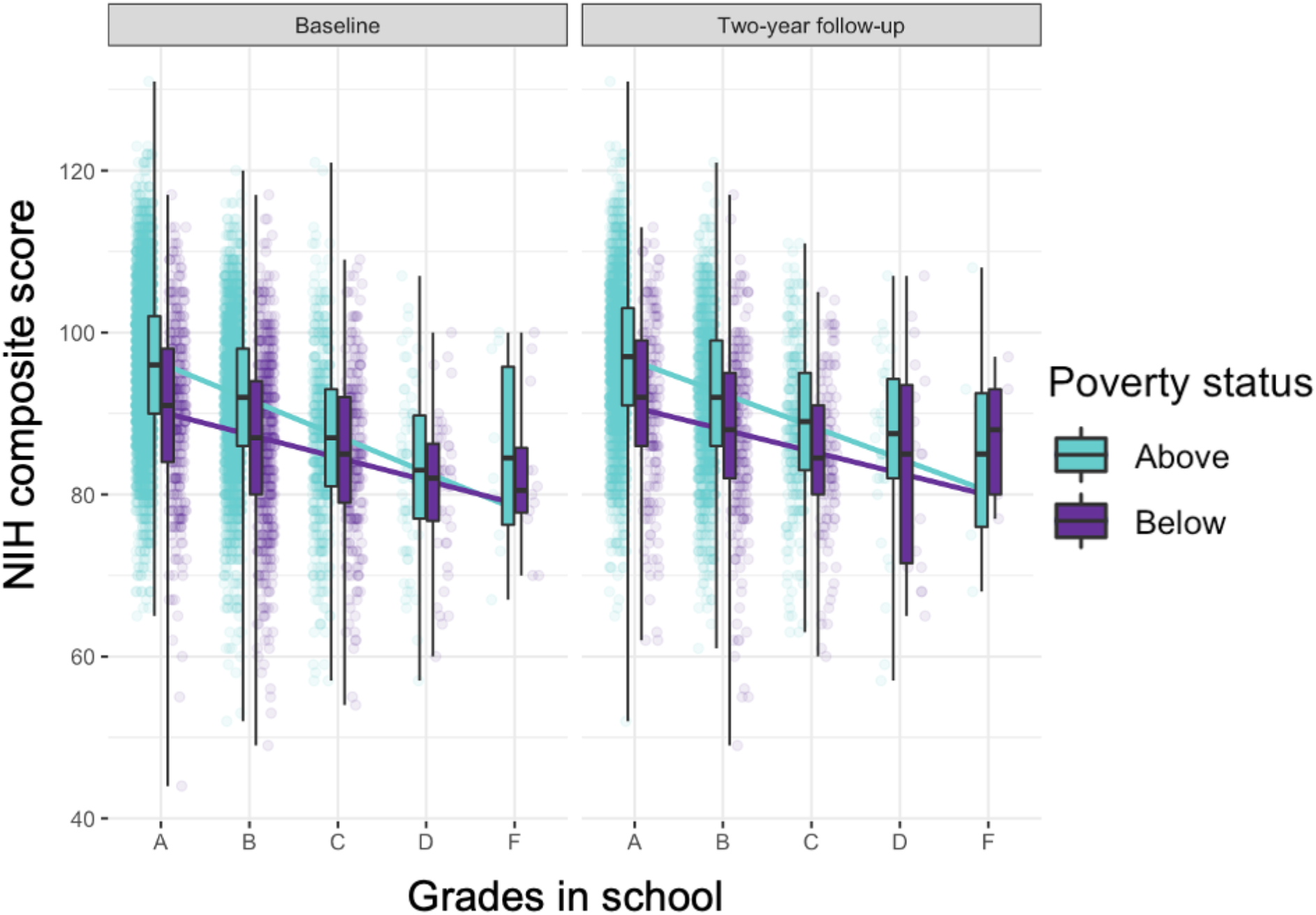
Relations between children’s test performance at baseline (T0) and their grades in school at baseline (T0; left panel) and two-year follow up (T2; right panel). Each point represents a different child; lighter teal color indicates children above poverty, while purple indicates children below poverty. Box plots for both groups at both timepoints are also displayed.

#### Longitudinal relations between test scores and grades

We expected to observe a similar pattern of results longitudinally – that is, when testing whether cognitive test performance at T0 predicts future academic performance. Higher cognitive test scores were related to higher grades in school one year later, controlling for grades at T0, *B* = -0.04, SD = 0.005, χ^2^ (2) = 72.81, *p* <.001. Contrary to predictions, this relation did not interact significantly as a function of poverty status, interaction: *B* = 0.02, SD = 0.01, χ^2^ (1) = 2.79, *p* = .095 (Figure 1). Thus, we found that cognitive test performance is somewhat predictive of concurrent and future academic performance, consistent with prior work. This relation was stronger for children above poverty at T0, but not longitudinally after controlling for T0.

Because grades at the two timepoints were correlated (*r* = .60; see Supplementary Figure 2), we conducted the same longitudinal analysis at T2 without controlling for grades at T0. This exploratory analysis revealed a significant interaction between children’s cognitive test performance at T0 and poverty status in predicting grades at T2 (interaction at T2: χ^2^ (1) = 9.55, *p* = 0.002). Mirroring results at T0, follow-up analyses showed a significant relation at T2 both for children above and below poverty, though the effect was stronger for children above poverty (below poverty: *B* = -0.04, SD = 0.01, χ^2^ (1) = 26.55, *p* < .001; above poverty: *B* = -0.07, SD = 0.004, χ^2^ (1) = 291.74, *p* < .001). Thus, grades at both timepoints were more strongly correlated with baseline cognitive test performance for children above than below poverty.

### Aim 3. Investigating associations between network connectivity and children’s grades in school and attention problems

#### Relations between academic performance and network connectivity

##### Concurrent relations between grades and network connectivity

As primary analyses, we next asked whether the relation between LFPN-DMN connectivity and children’s academic performance differed as a function of poverty status. We predicted that higher LFPN-DMN connectivity would be associated with lower grades for children above poverty but higher grades for children in poverty. On average, higher LFPN-DMN connectivity was related to worse grades concurrently at T0, *B* = 1.17, SD = 0.51, χ^2^ (2) = 9.23, *p* = .010. However, as predicted, this relation differed significantly as a function of poverty status, interaction: *B* = -3.11, SD = 1.11, χ^2^ (1) = 7.94, *p* = .005 (Figure 2). Follow-up analyses revealed that higher LFPN-DMN connectivity was related to worse grades for children above poverty; by contrast, it was directionally, though non-significantly, related to better grades for children below poverty (above poverty: *B* = 1.20, SD = 0.51, χ^2^ (1) = 5.51, *p* = .019; below poverty: *B* = -1.58, SD = 0.98, χ^2^ (1) = 2.61, *p* = .106). Thus, as predicted, LFPN-DMN connectivity was differentially associated with academic performance for children above and below poverty. Secondary analyses showed that, as predicted, higher CON-LFPN connectivity was also associated with worse grades concurrently, for children above and below poverty (see Supplement).

**Figure 2.**
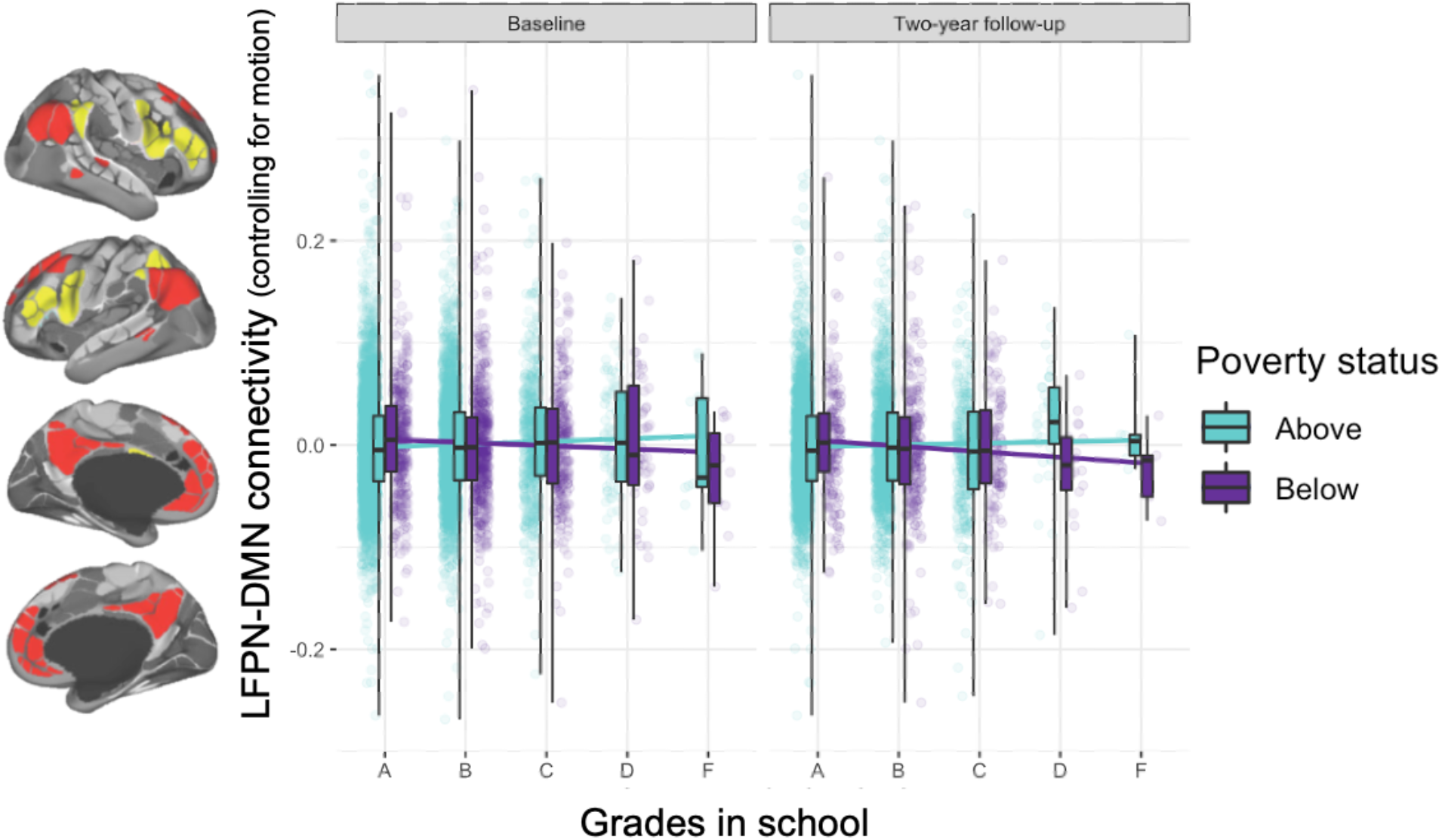
Relations between children’s LFPN-DMN connectivity at baseline (T0), after controlling for head motion, and their grades in school at baseline (T0; left panel) and two-year follow up (T2; right panel). Each point represents a different child; lighter teal color indicates children above poverty, while purple indicates children below poverty. Box plots for both groups at both timepoints are also displayed.

#### Longitudinal relations between grades and network connectivity

As primary analyses, we also conducted LFPN-DMN analyses longitudinally, to test the hypothesis that LFPN-DMN connectivity supports knowledge acquisition over the course of two years. We predicted that LFPN-DMN connectivity at T0 would be longitudinally associated with grades at T2. In contrast to this prediction, we found that there was no significant relation between LFPN-DMN connectivity and grades at T2 after controlling for T0 grades, *B* = 0.11, SD = 0.75, χ^2^ (2) = 0.88, *p* = .645, and this relation did not differ as a function of poverty status, interaction: *B* = -1.52, SD = 1.69, χ^2^ (1) = 0.81, *p* = .369.

Because grades at the two timepoints were correlated (*r* = .60; see Supplementary Figure 2), we conducted the same longitudinal analyses at T2 without controlling for grades at T0. This exploratory analysis revealed a significant interaction between children’s LFPN-DMN connectivity and poverty status at T0 in predicting grades at T2 (interaction at T2: χ^2^ (1) = 4.73, *p* = .030). Mirroring results at T0, the direction of the relation between LFPN-DMN connectivity and grades differed for the two groups, though it was not significant for either (below poverty: *B* = -2.50, SD = 1.36, χ^2^ (1) = 3.44, *p* = .064; above poverty: *B* = 0.49, SD = 0.70, χ^2^ (1) = 0.51, *p* = .475). Thus, the differential relation between connectivity at baseline and children’s grades was observed at two timepoints separated by two years.

We also tested associations longitudinally at an intermediate timepoint between T0 and T2 (T1), although we had not preregistered analyses involving T1 data. As reported in the Supplement, we found the expected interaction at T1, whereby higher LFPN-DMN connectivity appeared to be related to worse grades for children above poverty, but directionally related to better grades for children below poverty, even after controlling for grades at T0.

#### Relations between attention problems and network connectivity

As with grades, we next tested associations between children’s attention problems at T0 and T2, and network connectivity at T0. We predicted that stronger LFPN-DMN connectivity would be associated with greater attention problems longitudinally. Moreover, given our prior findings, we hypothesized that children below poverty might show the opposite pattern, such that higher connectivity would be related to fewer attention problems.

##### Concurrent relations between attention problems and network connectivity

While not pre-registered, we first tested cross-sectional associations to parallel analyses with grades and to set the stage for longitudinal analyses. On average, higher LFPN-DMN connectivity was related to more attention problems concurrently, *B* = 3.57, SD = 1.26, χ^2^ (2) = 10.33, *p* = .006; importantly, however, this relation differed significantly as a function of poverty status, interaction: *B* = -7.58, SD = 2.95, χ^2^ (1) = 6.61, *p* = .010 (Figure 3). While higher LFPN-DMN connectivity was related to more severe attention problems for children above poverty, it was not related to attention problems for children below poverty (above poverty: *B* = 3.72, SD = 1.20, χ^2^ (1) = 9.55, *p* = .002; below poverty: *B* = -3.70, SD = 3.33, χ^2^ (1) = 1.24, *p* = .265). Thus, stronger LFPN-DMN connectivity was associated with not only worse grades but also worse attention problems for children above poverty, but this was not the case for children below poverty.

**Figure 3.**
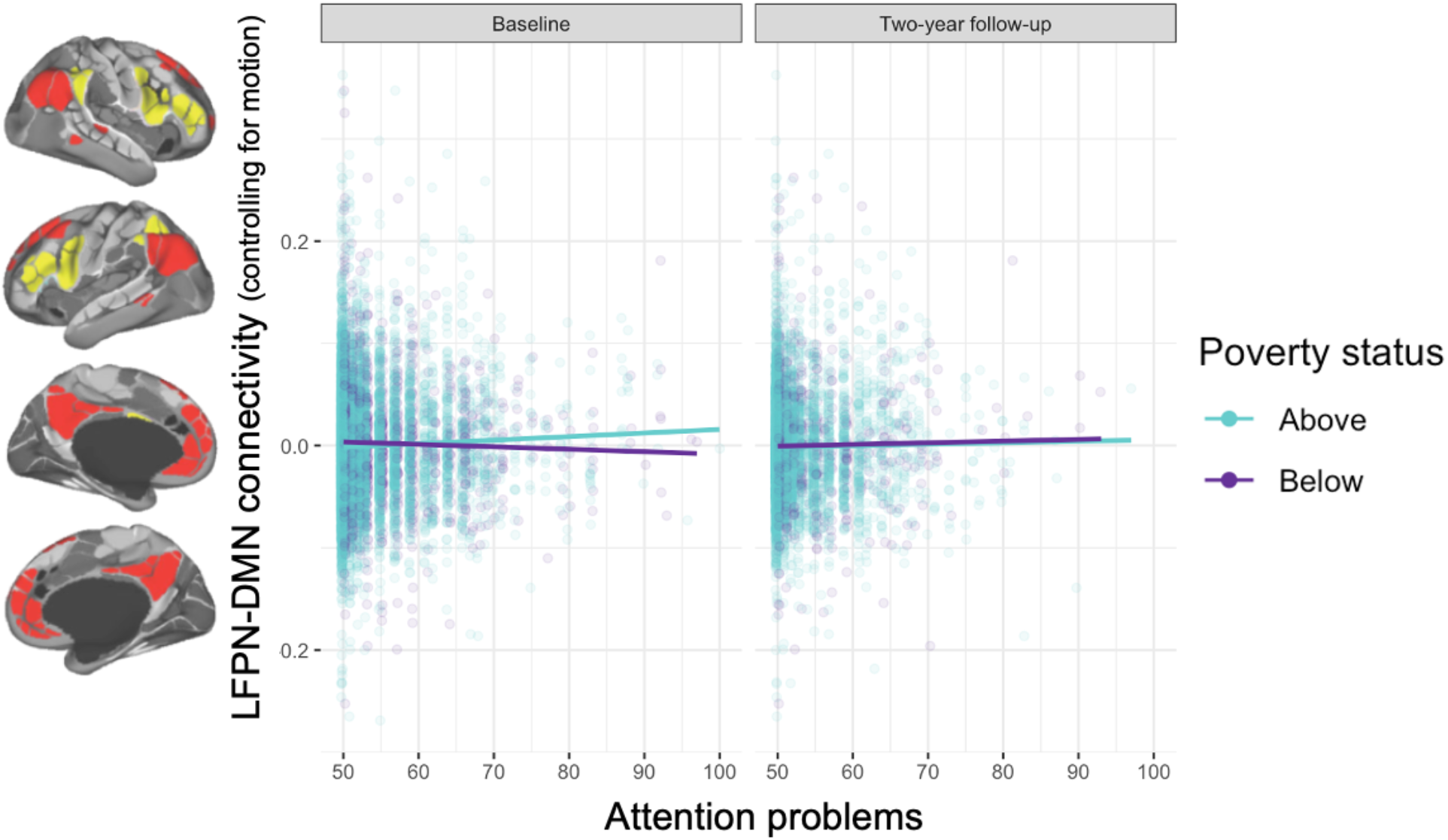
Relations between children’s LFPN-DMN connectivity at baseline (T0), after controlling for head motion, and their attention problems at baseline (T0; left panel) and two-year follow up (T2; right panel). Each point represents a different child; lighter teal color indicates children above poverty, while purple indicates children below poverty.

As secondary analyses, we also tested whether patterns of CON connectivity at baseline were differentially associated with attention problems for children above and below poverty, after accounting for LFPN-DMN connectivity and its interaction with poverty status. As predicted, we found that higher CON-DMN connectivity was associated with more attention problems, and this effect was strongest for children in poverty. Contrary to predictions, we did not find any significant associations with CON-LFPN connectivity.

##### Longitudinal relations between attention problems and network connectivity

We next tested our hypothesis that higher LFPN-DMN connectivity would be associated with more attention problems longitudinally, controlling for attention at T0. Contrary to our prediction, we found no significant relation between LFPN-DMN and attention at T2 when controlling for attention at T0 (T1: *B* = -0.89, SD = 0.84, χ^2^ (2) = 1.49, *p* = .474; T2: *B* = -0.14, SD = 1.12, χ^2^ (2) = 0.17, *p* = .919). Further, this relation did not differ significantly as a function of poverty status (interaction: *B* = 1.15, SD = 2.77, χ^2^ (1) = 0.17, *p* = .684; Figure 3).

Because attention problems were correlated across timepoints (*r* = .68; see Supplementary Figure 2), we also conducted an exploratory analysis without controlling for attention at T0. There was similarly no interaction at T2 when not controlling for attention at T0, *X*^2^ (1) = 0.010, *p* = .925. Thus, individual variability in attention problems was linked to LFPN-DMN connectivity only at ages 9-10—and at that time, it was linked in opposite directions as a function of poverty status.

#### Summary of results

A summary of our pre-registered hypotheses, analyses, and results can be found in Supplementary Table 1. Overall, we found that CON-DMN and CON-LFPN connectivity declined over early adolescence, but LFPN-DMN connectivity did not. Children above poverty showed a steeper decline in CON-DMN connectivity than those below. Turning to behavioral measures, we found that children’s scores on standardized cognitive tests were related to their grades in school both concurrently and longitudinally, though the association was attenuated for children in poverty. Finally, we found all three networks were associated with children’s grades and attention problems. Confirming hypotheses, the link between LFPN-DMN connectivity and behavior differed for children above and below poverty, with lower LFPN-DMN connectivity being adaptive only for children above poverty. On the other hand, lower CON-LFPN connectivity was uniformly related to better grades, while lower CON-DMN was related to more attention problems, particularly among children in poverty.

## Discussion

In this study, we sought to investigate trajectories of LFPN and DMN, as well as CON, network coupling over middle childhood and early adolescence, and their relation to academic and behavioral resilience. To this end, we examined rs-fMRI network coupling longitudinally in relation to grades and attention problems in a diverse sample of participants at two timepoints approximately two years apart, spanning ages 9-13 across the sample. The central goal of this study was to assess whether associations between functional connectivity and performance differ meaningfully between children whose families lived above and below poverty. Unlike studies that have examined brain differences between relatively higher- and lower-income children, this study included children living below the poverty line, some of whom are able to cope with severe adversity. Further, by examining more ecologically valid measures of children’s cognitive performance, we sought to better capture the neural basis of children’s resilience with regard to societal constraints.

On the cognitive side, we found that there is not a universally strong link between performance on tests of executive functioning and these more real-world outcome measures. Despite countless studies showing relations between performance on these tests and performance in school (e.g., Best et al., 2011; Cowan, 2014; St Clair-Thompson & Gathercole, 2006; Willoughby et al., 2019), we found that cognitive test scores were more highly correlated with academic performance in children above than below poverty. The lower correlation for children below poverty likely reflects obstacles that hinder their ability to reach their full potential, such as fewer school resources and lower quality of instruction (Horng, 2005; Orfield & Lee, 2005; Reardon & Owens, 2014) and discrimination at school (Darling-Hammond, 2001; Hettleman, 2003; Scott et al., 2020).

The weaker link between test scores and school performance for children below poverty could stem in part from the use of cognitive tests that are argued to be culturally biased methods of assessment (Miller-Cotto et al., 2021). Of course, scholastic assessments are themselves considered to be culturally biased—a concern that led to the development of cognitive tests designed to measure aptitude without requiring extensive background knowledge. Most importantly, for our purposes, academic performance is directly tied to children’s longer-term prospects, whereas performance on abstract cognitive tests is not. The group difference in the strength of the association between these metrics highlights the importance of examining real-world outcomes in relation to brain network development.

### Network associations with children’s grades

Turning to associations with children’s resting state connectivity, we found that having lower LFPN-DMN connectivity was concurrently related to having better grades—but only among the children above poverty; for children living below poverty, this association was in the opposite direction. An exploratory analysis found that the same dissociation was present for grades two years later; however, because grades at T2 and T0 were correlated, this relation was not significant when controlling for grades at T0. Additional exploratory analyses suggested that the dissociation was significant at T1, even when controlling for grades at T0, pointing to its relative stability over time.

Further, the data did not suggest that high-performing children below poverty exhibited a pattern of connectivity that was beneficial at age 9-10 but that later became detrimental. Instead, there are qualitative differences in what appears to be academically adaptive for children above and below poverty; these differences may be established earlier in childhood and then attenuate slightly across middle childhood and early adolescence.

We also found that higher CON-LFPN was related to worse academic performance for both children above and below poverty. By contrast, prior evidence suggests that higher CON between-network connectivity is related to better executive functioning (Marek et al., 2015). Further, though there is little work linking children’s performance in school to their resting state connectivity, one notable study found that greater network integration at ages 7-9 was associated with higher scholastic performance (Chaddock-Heyman et al., 2018). As there are a number of possible reasons for these discrepant findings, including differences in age range, additional work will be needed to reconcile them.

### Network associations with children’s attention problems

With regard to attention problems, our primary hypotheses focused on longitudinal associations, given a prior study showing that LFPN-DMN connectivity at age 7 predicted changes in attention problems across four years, controlling for attention problems at the initial timepoint (Whitfield-Gabrieli et al., 2020). Here, we did not find that LFPN-DMN connectivity at baseline was related to attention problems two years later, either for children above or below poverty, when controlling for baseline attention problems. There are several possible reasons for our discrepant findings with the prior study (Whitfield-Gabrieli et al., 2020), including its seed-based approach, longer lag between timepoints, younger ages at baseline, and small sample size.

However, exploratory analyses examining cross-sectional associations found results broadly consistent with prior evidence: stronger LFPN-DMN connectivity was associated with greater attention problems for children above poverty. For children below poverty, on the other hand, this effect was non-significant and in the opposite direction, and the group interaction was significant. In contrast to our findings with grades, this interaction was not evident longitudinally, even when not controlling for baseline attention, suggesting that the dissociation between LFPN-DMN connectivity and attention problems for children above and below poverty is attenuated with age.

Importantly, other network interactions appear to be more reliable markers of attention problems for children below poverty than is LFPN-DMN. Indeed, secondary and exploratory analyses in the present study indicate that stronger CON-DMN connectivity was associated with more severe attention problems for children below poverty, both concurrently and longitudinally. Additionally, another study involving the ABCD dataset showed that weaker anti-correlations between the Dorsal Attention Network and the DMN is associated with more attention problems, and that it is also related to socioeconomic status (Owens et al., 2020). Thus, segregation of the DMN from both the CON and Dorsal Attention Network may be associated with better behavioral outcomes, for children above and below poverty.

### Network changes over time

Based on prior literature, we anticipated that the LFPN and DMN would become less coupled over time across the full sample (Baum et al., 2017; Grayson & Fair, 2017; Sherman et al., 2014). However, we found that LFPN-DMN connectivity did not change consistently over the two-year study period; connectivity decreased for some individuals and increased for others. This null result may reflect the relatively brief time window (two years) and/or the particular age range over which we examined changes (9-11 at the first timepoint; 10-13 at the third). We note that one prior study found increasing segregation between nodes of the DMN and LFPN longitudinally over ages 10-11, though this research used a seed-based approach that differed from our network-based approach (Sherman et al., 2014).

Thus, there was no consistent developmental change in LFPN-DMN connectivity across the two years for either group. However, because we had seen differential patterns of connectivity across poverty levels as a function of cognitive performance, we additionally conducted exploratory analyses testing whether children’s cognitive test scores at baseline influenced individuals’ trajectory of change in connectivity. However, this possibility was not borne out by the data either: trajectories of LFPN-DMN connectivity did not differ as a function of children’s initial cognitive test scores.

Other studies have provided evidence of accelerated physical and brain development among children growing up in adversity, which could help them to adapt more readily to harsher living conditions (Belsky, 2019; Callaghan & Tottenham, 2016; Gee et al., 2013; McDermott et al., 2021; Tooley et al., 2021). Here, we do not see evidence for accelerated rate of LFPN-DMN network development for children below poverty—at least, not at this point in development. However, it is possible that a differential trajectory of change is visible earlier in childhood, at a time when networks affiliations are changing more markedly and/or more consistently.

Given prior literature indicating that the LFPN and DMN interact with the CON, a network that has also been implicated in cognitive functioning, we also explored the development of CON connectivity. We found that CON-LFPN and CON-DMN connectivity decreased longitudinally, on average. These findings fit with the broad characterization of network segregation over development, but differ from past studies showing increased CON integration with other brain networks from ages 8-21 (Lopez et al., 2020) or 10-26 (Marek et al., 2015). This discrepancy could stem from the fact that our participants were at the youngest end of the broad age ranges reported in these cross-sectional studies; it is also possible that two years was insufficiently long to see significant change. Additionally or alternatively, discrepancies in results could be related to the different connectivity metrics used across studies.

Importantly, however, we also found that the trajectory of change in between-network connectivity for the CON differed as a function of whether children were in poverty at baseline. This interaction was significant for CON-DMN connectivity but not CON-LFPN connectivity. Follow-up analyses revealed that only the children above poverty showed a significant decrease in CON-DMN connectivity. There was no significant change for children below poverty—and we showed subsequently that this did not depend on cognitive performance. This pattern of results fits with findings of other recent work (Chahal et al., 2022) and raises the possibility that the children living below poverty reached CON network maturity slightly earlier, potentially consistent with the accelerated development hypothesis for these particular networks (Belsky, 2019; Callaghan & Tottenham, 2016; Gee et al., 2013; McDermott et al., 2021; Tooley et al., 2021). To ascertain which neural systems show accelerated maturation in the face of adversity, it will be necessary to test for differential trajectories during a dynamic window of development for a given system.

Of note, exploratory analyses suggested the rate of network change for LFPN-DMN, CON-DMN, and CON-LFPN did not interact with children’s initial cognitive test performance. Thus, there is no evidence that there is any one trajectory of network change that is particularly adaptive in terms of cognitive performance, either for children above or below poverty. We therefore cannot conclude that children in poverty who show more evidence for accelerated CON maturation than their peers are at greater risk in terms of cognitive development. Clearly, more research is needed that continues to follow these trajectories—and their associations with later outcomes—over time.

### Conclusions, limitations, and future directions

Overall, these results build on a prior study suggesting that lower LFPN-DMN connectivity is adaptive for higher-but not lower-income children, as measured by performance on tests of executive functioning and reasoning (Ellwood-Lowe et al., 2021). Here, we extend this result in several ways. In particular, we show the same pattern for more ecologically valid measures that capture children’s resilience in real-world contexts and—given the importance of scholastic achievement for upward social mobility—that can directly impact their opportunities in life. Further, we show that the dissociation observed previously is, at the very least, not linked to worse outcomes over the longer term for higher-performing children in poverty.

The phenomena established by these initial results across two studies lay a foundation for more detailed analysis of functional connectivity. For example, it may be useful to explore subnetworks of LFPN and DMN, given distinctions in their contributions to cognition (Buckner & DiNicola, 2019; Dixon et al., 2018; Fornito et al., 2012; Lopez et al., 2020). In addition, it will be important to assess individual-level networks (Seitzman et al., 2019), to see whether network boundaries differ meaningfully as a function of children’s experiences. Further, it would be interesting to determine which specific aspects of children’s home environment underlie the effects reported here, given that experiences differ markedly even among children living in poverty (DeJoseph et al., 2021; Ellwood-Lowe et al., 2021; Humphreys & Zeanah, 2015; McLaughlin et al., 2014; Rakesh, Seguin, et al., 2021; Rakesh, Zalesky, et al., 2021).

While the differential patterns of brain activity we see here may reflect years of childhood experiences, other research has shown that rs-fMRI changes as a function of even brief experiences, even in adulthood (Guerra-Carrillo et al., 2014). Similarly, the developmental trajectory of brain coupling is likely not immutable; to the extent that an individual brain show stability in connectivity over time, this could in large part reflect stability in the context in which they live—their challenges and opportunities.

With the current data, we cannot say whether these dissociations in rs-fMRI connectivity would also be reflected in children’s performance during externally-directed cognitive tasks, or whether they simply represent cumulative differences in other thought patterns. In other words, do high-performing children in poverty deactivate DMN less during cognitive task performance? Or do they engage in more creative thinking in day-to-day life, for example, or other thought patterns that frequently coactivate these networks? Future studies should measure LFPN and DMN coupling during cognitive task performance and whether it differs as a function of poverty status. Of note for future investigations, other networks may also show differential relations with behavior as a function of poverty status.

Given the societal relevance of this work, a cautionary note is warranted. The effect sizes were quite small, and there was substantial overlap in network connectivity and its relation to behavior between children living above and below poverty. Because children below poverty are typically underrepresented in neuroimaging research, we chose to examine them as a separate group, defined based on their combined family income and the number of people in their household (see also Ellwood-Lowe et al., 2021). Of course, this is a somewhat arbitrary distinction based on an estimate of whether a child’s family has the financial resources they need to meet their basic needs; more than likely, this dataset includes a substantial number of children in poverty who have more common experiences with those above poverty, and vice versa. Numerous experiences beyond financial resources shape mental processes. In addition, numerous other features of brain structure and function contribute to these individual differences in mental processing. Further, we only considered poverty status at baseline, but future studies should explore whether effects change for children who move in or out of poverty over adolescence.

Compounding the issue of underrepresentation of certain groups in research, the children who were included in our sample differed meaningfully from those who were excluded based on low fMRI data quality or incomplete behavioral data. On average, children who were excluded were more likely to be in poverty; they were also more likely to be younger, be male, have more attention problems, have lower grades, and have lower cognitive test scores. Across the socioeconomic spectrum, demands associated with the study—in particular, producing usable rs-fMRI data—selected for children who were older, female, and higher-performing. Thus, we cannot say whether the same patterns as in our current study would be found for children who were among the lowest performing in both the below and above poverty groups. More broadly, this differential exclusion illustrates how even representative neuroimaging studies lose important information about children who struggle to stay still in the scanner, contributing to difficulties making accurate generalizations about children (Falk et al., 2013). Despite these issues, we should note that our remaining sample is still quite diverse with respect to all of these characteristics.

Our findings highlight that a one-size-fits-all approach to promoting healthy brain development may not be possible, given the inequities of structural barriers faced by different students. These patterns of resting-state functional connectivity suggest that students in poverty who perform well may rely on different thought patterns to do so. Moreover, this short-term cognitive resilience may come with more long-term costs for children in poverty. For example, the burden of trying to adapt to unfair structural conditions may contribute to chronic stress and increased allostatic load (McEwen & Wingfield, 2003); future research should investigate this possibility. Indeed, there is recent evidence from the ABCD sample that patterns of brain development are associated meaningfully with the implementation of social and economic policies aimed at mitigating resource strain for low SES children, pointing to the importance of focusing efforts on removing structural barriers to children’s success (Weissman et al., 2021).

Taken together, these results show that the cognitive and neural factors that influence achievement are not exactly the same for children above and below poverty. Within a deficit framework, a goal toward promoting equity in academic achievement might be to “correct” brain networks, such that children below poverty show a pattern more closely resembling that of children above poverty. The findings presented here complicate this idea, suggesting that in the absence of taking children out of poverty, approaches that maximize their specific developmental trajectories and capacities may be needed. Our findings also highlight the importance of recruiting diverse samples for understanding human development; even among children living within the United States, who themselves share many experiences in common, there appear to be important experience-dependent differences in patterns of brain network development that support academic and behavioral resilience.

## Acknowledgements

This study would not be possible without the massive efforts of the large team of ABCD leaders and organizers, staff and data curators, and families and children who participated. We are grateful to members of the Building Blocks of Cognition Lab for their support. The content is solely the responsibility of the authors and does not necessarily represent the official views of the National Institutes of Health. MEL was supported by NSF GRFP DGE 1752814. Data used in the preparation of this article were obtained from the Adolescent Brain Cognitive Development^SM^ (ABCD) Study (https://abcdstudy.org), held in the NIMH Data Archive (NDA). This is a multisite, longitudinal study designed to recruit more than 10,000 children age 9-10 and follow them over 10 years into early adulthood. The ABCD Study® is supported by the National Institutes of Health and additional federal partners under award numbers U01DA041048, U01DA050989, U01DA051016, U01DA041022, U01DA051018, U01DA051037, U01DA050987, U01DA041174, U01DA041106, U01DA041117, U01DA041028, U01DA041134, U01DA050988, U01DA051039, U01DA041156, U01DA041025, U01DA041120, U01DA051038, U01DA041148, U01DA041093, U01DA041089, U24DA041123, U24DA041147. A full list of supporters is available at https://abcdstudy.org/federal-partners.html. A listing of participating sites and a complete listing of the study investigators can be found at https://abcdstudy.org/consortium_members/. ABCD consortium investigators designed and implemented the study and/or provided data but did not necessarily participate in the analysis or writing of this report. This manuscript reflects the views of the authors and may not reflect the opinions or views of the NIH or ABCD consortium investigators. The ABCD data repository grows and changes over time. The ABCD data used in this report came from the fast track data release. The raw data are available at https://nda.nih.gov/edit_collection.html?id=2573. Instructions on how to create an NDA study are available at https://nda.nih.gov/training/modules/study.html).

## References

Achenbach, T. M., & Ruffle, T. M. (2000). The child behavior checklist and related forms for assessing behavioral/emotional problems and competencies. Pediatrics in Review, 21(8), 265–271. https://doi.org/10.1542/pir.21-8-265

Agnoli, S., Vanucci, M., Pelagatti, C., & Corazza, G. E. (2018). Exploring the Link Between Mind Wandering, Mindfulness, and Creativity: A Multidimensional Approach. Creativity Research Journal, 30(1), 41–53. https://doi.org/10.1080/10400419.2018.1411423

Barch, D. M., Albaugh, M. D., Avenevoli, S., Chang, L., Clark, D. B., Glantz, M. D., Hudziak, J. J., Jernigan, T. L., Tapert, S. F., Yurgelun-Todd, D., Alia-Klein, N., Potter, A. S., Paulus, M. P., Prouty, D., Zucker, R. A., & Sher, K. J. (2018). Demographic, physical and mental health assessments in the adolescent brain and cognitive development study: Rationale and description. Developmental Cognitive Neuroscience, 32(March 2017), 55–66. https://doi.org/10.1016/j.dcn.2017.10.010

Bates, D., Mächler, M., Bolker, B., & Walker, S. (2015). Fitting Linear Mixed-Effects Models Using {lme4}. Journal of Statistical Software, 67(1), 1–48. https://doi.org/10.18637/jss.v067.i01

Baum, G. L., Ciric, R., Roalf, D. R., Betzel, R. F., Moore, T. M., Shinohara, R. T., Kahn, A. E., Vandekar, S. N., Rupert, P. E., & Quarmley, M. (2017). Modular segregation of structural brain networks supports the development of executive function in youth. Current Biology, 27(11), 1561–1572.

Beaty, R. E., Benedek, M., Silvia, P. J., & Schacter, D. L. (2016). Creative Cognition and Brain Network Dynamics. Trends in Cognitive Sciences, 20(2), 87–95. https://doi.org/10.1016/j.tics.2015.10.004

Beaty, R. E., Silvia, P. J., & Benedek, M. (2017). Brain networks underlying novel metaphor production. Brain and Cognition, 111, 163–170. https://doi.org/10.1016/j.bandc.2016.12.004

Belsky, J. (2019). Early-life adversity accelerates child and adolescent development. Current Directions in Psychological Science, 28(3), 241–246.

Best, J. R., Miller, P. H., & Naglieri, J. A. (2011). Relations between executive function and academic achievement from ages 5 to 17 in a large, representative national sample. Learning and Individual Differences, 21(4), 327–336.

Buckner, R. L., & DiNicola, L. M. (2019). The brain’s default network: updated anatomy, physiology and evolving insights. In Nature reviews. Neuroscience (Vol. 20, Issue 10, pp. 593–608). NLM (Medline). https://doi.org/10.1038/s41583-019-0212-7

Callaghan, B. L., & Tottenham, N. (2016). The Stress Acceleration Hypothesis: Effects of early-life adversity on emotion circuits and behavior. Current Opinion in Behavioral Sciences, 7, 76–81. https://doi.org/10.1016/j.cobeha.2015.11.018

Casey, B. J., Cannonier, T., Conley, M. I., Cohen, A. O., Barch, D. M., Heitzeg, M. M., Soules, M. E., Teslovich, T., Dellarco, D. V., Garavan, H., Orr, C. A., Wager, T. D., Banich, M. T., Speer, N. K., Sutherland, M. T., Riedel, M. C., Dick, A. S., Bjork, J. M., Thomas, K. M., … Dale, A. M. (2018). The Adolescent Brain Cognitive Development (ABCD) study: Imaging acquisition across 21 sites. Developmental Cognitive Neuroscience, 32(January), 43–54. https://doi.org/10.1016/j.dcn.2018.03.001

Chaddock-Heyman, L., Weng, T. B., Kienzler, C., Erickson, K. I., Voss, M. W., Drollette, E. S., Raine, L. B., Kao, S. C., Hillman, C. H., & Kramer, A. F. (2018). Scholastic performance and functional connectivity of brain networks in children. PLoS ONE, 13(1), 1–16. https://doi.org/10.1371/journal.pone.0190073

Chahal, R., Miller, J. G., Yuan, J. P., Buthmann, J. L., & Gotlib, I. H. (2022). An exploration of dimensions of early adversity and the development of functional brain network connectivity during adolescence: Implications for trajectories of internalizing symptoms. Development and Psychopathology, 1–15.

Chai, X. J., Ofen, N., Gabrieli, J. D. E., & Whitfield-Gabrieli, S. (2014). Selective development of anticorrelated networks in the intrinsic functional organization of the human brain. Journal of Cognitive Neuroscience. https://doi.org/10.1162/jocn_a_00517

Christensen, R. H. B. (2018). Cumulative link models for ordinal regression with the R package ordinal. Submitted in J. Stat. Software.

Christoff, K., Gordon, A. M., Smallwood, J., Smith, R., & Schooler, J. W. (2009). Experience sampling during fMRI reveals default network and executive system contributions to mind wandering. Proceedings of the National Academy of Sciences of the United States of America, 106(21), 8719–8724. https://doi.org/10.1073/pnas.0900234106

Cowan, N. (2014). Working memory underpins cognitive development, learning, and education. Educational Psychology Review, 26(2), 197–223.

Darling-Hammond, L. (2001). Inequality in teaching and schooling: How opportunity is rationed to students of color in America. BD Smedley, AY Stith, L. Colburn C. & H. Evans (Eds.), The Right Thing to Do—The Smart Thing to Do, 208–233.

DeFelipe, J. (2006). Brain plasticity and mental processes: Cajal again. Nature Reviews Neuroscience, 7(10), 811–817. https://doi.org/10.1038/nrn2005

DeJoseph, M. L., Sifre, R. D., Raver, C. C., Blair, C. B., & Berry, D. (2021). Capturing Environmental Dimensions of Adversity and Resources in the Context of Poverty Across Infancy Through Early Adolescence: A Moderated Nonlinear Factor Model. Child Development. https://doi.org/10.1111/cdev.13504

DeSerisy, M., Ramphal, B., Pagliaccio, D., Raffanello, E., Tau, G., Marsh, R., Posner, J. & Margolis, A. E. (2021). Frontoparietal and default mode network connectivity varies with age and intelligence. Developmental cognitive neuroscience, 48, 100928.

Desikan, R. S., Segonne, F., Fischl, B., Quinn, B. T., Dickerson, B. C., Blacker, D., Buckner, R. L., Dale, A. M., Maguire, R. P., Hyman, B. T., Albert, M. S., & Killiany, R. J. (2006). An automated labeling system for subdividing the human cerebral cortex on MRI scans into gyral based regions of interest. In NeuroImage.

Destrieux, C., Fischl, B., Dale, A., & Halgren, E. (2010). Automatic parcellation of human cortical gyri and sulci using standard anatomical nomenclature. Neuroimage, 53(1), 1–15.

Diamond, M. C., Krech, D., & Rosenzweig, M. R. (1964). The effects of an enriched environment on the histology of the rat cerebral cortex. Journal of Comparative Neurology, 123(1), 111–119. https://doi.org/10.1002/cne.901230110

Dixon, M. L., De La Vega, A., Mills, C., Andrews-Hanna, J., Spreng, R. N., Cole, M. W., & Christoff, K. (2018). Heterogeneity within the frontoparietal control network and its relationship to the default and dorsal attention networks. Proceedings of the National Academy of Sciences of the United States of America, 115(13), E3068. https://doi.org/10.1073/pnas.1803276115

Dixon, M. L., Fox, K. C. R., & Christoff, K. (2014). A framework for understanding the relationship between externally and internally directed cognition. Neuropsychologia, 62, 321–330. https://doi.org/10.1016/j.neuropsychologia.2014.05.024

Dosenbach, N. U. F., Koller, J. M., Earl, E. A., Miranda-Dominguez, O., Klein, R. L., Van, A. N., Snyder, A. Z., Nagel, B. J., Nigg, J. T., Nguyen, A. L., Wesevich, V., Greene, D. J., & Fair, D. A. (2017). Real-time motion analytics during brain MRI improve data quality and reduce costs. NeuroImage, 161, 80–93. https://doi.org/https://doi.org/10.1016/j.neuroimage.2017.08.025

Ellwood-Lowe, M. E., Whitfield-Gabrieli, S., & Bunge, S. A. (2021). Brain network coupling associated with cognitive performance varies as a function of a child’s environment in the ABCD study. Nature Communications, 12(1), 1–14. https://doi.org/10.1038/s41467-021-27336-y

Falk, E. B., Hyde, L. W., Mitchell, C., Faul, J., Gonzalez, R., Heitzeg, M. M., … & Schulenberg, J. (2013). What is a representative brain? Neuroscience meets population science. Proceedings of the National Academy of Sciences, 110(44), 17615–17622.

Finn, A. S., Minas, J. E., Leonard, J. A., Mackey, A. P., Salvatore, J., Goetz, C., West, M. R., Gabrieli, C. F. O., & Gabrieli, J. D. E. (2017). Functional brain organization of working memory in adolescents varies in relation to family income and academic achievement. Developmental Science, 20(5). https://doi.org/10.1111/desc.12450

Fornito, A., Harrison, B. J., Zalesky, A., & Simons, J. S. (2012). Competitive and cooperative dynamics of large-scale brain functional networks supporting recollection. Proceedings of the National Academy of Sciences of the United States of America, 109(31), 12788–12793. https://doi.org/10.1073/pnas.1204185109

Friston, K. J., Ashburner, J., Frith, C. D., Poline, J.-B., Heather, J. D., & Frackowiak, R. S. J. (1995). Spatial registration and normalization of images. Human Brain Mapping, 3(3), 165–189. https://doi.org/https://doi.org/10.1002/hbm.460030303

Garavan, H., Bartsch, H., Conway, K., Decastro, A., Goldstein, R. Z., Heeringa, S., Jernigan, T., Potter, A., Thompson, W., & Zahs, D. (2018). Recruiting the ABCD sample: Design considerations and procedures. Developmental Cognitive Neuroscience, 32(April), 16–22. https://doi.org/10.1016/j.dcn.2018.04.004

Gee, D. G., Gabard-Durnam, L. J., Flannery, J., Goff, B., Humphreys, K. L., Telzer, E. H., Hare, T. A., Bookheimer, S. Y., & Tottenham, N. (2013). Early developmental emergence of human amygdala–prefrontal connectivity after maternal deprivation. Proceedings of the National Academy of Sciences, 110(39), 15638–15643.

Gordon, E. M., Laumann, T. O., Adeyemo, B., Huckins, J. F., Kelley, W. M., & Petersen, S. E. (2016). Generation and Evaluation of a Cortical Area Parcellation from Resting-State Correlations. Cerebral Cortex, 26(1), 288–303. https://doi.org/10.1093/cercor/bhu239

Grayson, D. S., & Fair, D. A. (2017). Development of large-scale functional networks from birth to adulthood: a guide to neuroimaging literature. NeuroImage. https://doi.org/10.1016/j.neuroimage.2017.01.079

Guerra-Carrillo, B., Mackey, A. P., & Bunge, S. A. (2014). Resting-state fMRI: A window into human brain plasticity. In Neuroscientist (Vol. 20, Issue 5, pp. 522–533). https://doi.org/10.1177/1073858414524442

Hagler, D. J., Hatton, S. N., Cornejo, M. D., Makowski, C., Fair, D. A., Dick, A. S., Sutherland, M. T., Casey, B. J., Barch, D. M., Harms, M. P., Watts, R., Bjork, J. M., Garavan, H. P., Hilmer, L., Pung, C. J., Sicat, C. S., Kuperman, J., Bartsch, H., Xue, F., … Dale, A. M. (2019). Image processing and analysis methods for the Adolescent Brain Cognitive Development Study. NeuroImage. https://doi.org/10.1016/j.neuroimage.2019.116091

Harpin, V. A. (2005). The effect of ADHD on the life of an individual, their family, and community from preschool to adult life. Archives of Disease in Childhood, 90(SUPPL. 1), 2–7. https://doi.org/10.1136/adc.2004.059006

Hettleman, K. R. (2003). The Invisible Dyslexics: How Public School Systems in Baltimore and Elsewhere Discriminate against Poor Children in the Diagnosis and Treatment of Early Reading Difficulties.

Hilger, K., Ekman, M., Fiebach, C. J., & Basten, U. (2017). Intelligence is associated with the modular structure of intrinsic brain networks. Scientific Reports, 7(1), 1–12. https://doi.org/10.1038/s41598-017-15795-7

Horng, E. L. (2005). Poor Working Conditions Make Urban Schools Hard-to-Staff.

Humphreys, K. L., & Zeanah, C. H. (2015). Deviations from the Expectable Environment in Early Childhood and Emerging Psychopathology. Neuropsychopharmacology : Official Publication of the American College of Neuropsychopharmacology, 40(April), 1–59. https://doi.org/10.1038/npp.2014.165

Jaarsveld, S., & Lachmann, T. (2017). Intelligence and creativity in problem solving: The importance of test features in cognition research. Frontiers in Psychology, 8(FEB), 1–12. https://doi.org/10.3389/fpsyg.2017.00134

Kucyi, A., Esterman, M., Capella, J., Green, A., Uchida, M., Biederman, J., Gabrieli, J. D. E., Valera, E. M., & Whitfield-Gabrieli, S. (2021). Prediction of stimulus-independent and task-unrelated thought from functional brain networks. Nature Communications, 12(1). https://doi.org/10.1038/s41467-021-22027-0

Leonard, J. A., Romeo, R. R., Park, A. T., Takada, M. E., Robinson, S. T., Grotzinger, H., Last, B. S., Finn, A. S., Gabrieli, J. D. E., & Mackey, A. P. (2019). Associations between cortical thickness and reasoning differ by socioeconomic status in development. Developmental Cognitive Neuroscience, 36(March 2018), 100641. https://doi.org/10.1016/j.dcn.2019.100641

Liston, C., McEwen, B. S., & Casey, B. J. (2009). Psychosocial stress reversibly disrupts prefrontal processing and attentional control. Proceedings of the National Academy of Sciences of the United States of America, 106(3), 912–917. https://doi.org/10.1073/pnas.0807041106

Lopez, K. C., Kandala, S., Marek, S., & Barch, D. M. (2020). Development of Network Topology and Functional Connectivity of the Prefrontal Cortex. Cerebral Cortex, 30(4), 2489–2505. https://doi.org/10.1093/cercor/bhz255

Luciana, M., Bjork, J. M., Nagel, B. J., Barch, D. M., Gonzalez, R., Nixon, S. J., & Banich, M. T. (2018). Adolescent neurocognitive development and impacts of substance use: Overview of the adolescent brain cognitive development (ABCD) baseline neurocognition battery. Developmental Cognitive Neuroscience, 32(February), 67–79. https://doi.org/10.1016/j.dcn.2018.02.006

Mackey, A. P., Singley, A. T. M., & Bunge, S. A. (2013). Intensive reasoning training alters patterns of brain connectivity at rest. Journal of Neuroscience, 33(11), 4796–4803. https://doi.org/10.1523/JNEUROSCI.4141-12.2013

Marek, S., Hwang, K., Foran, W., Hallquist, M. N., & Luna, B. (2015). The Contribution of Network Organization and Integration to the Development of Cognitive Control. PLoS Biology, 13(12), 1–25. https://doi.org/10.1371/journal.pbio.1002328

McDermott, C. L., Hilton, K., Park, A. T., Tooley, U. A., Boroshok, A. L., Mupparapu, M., Scott, J. A. M., Bumann, E. E., & Mackey, A. P. (2021). Early life stress is associated with earlier emergence of permanent molars. Proceedings of the National Academy of Sciences of the United States of America, 118(24), 3–5. https://doi.org/10.1073/pnas.2105304118

McEwen, B. S., & Wingfield, J. C. (2003). The concept of allostasis in biology and biomedicine. Hormones and Behavior, 43(1), 2–15. https://doi.org/10.1016/S0018-506X(02)00024-7

McLaughlin, K. A., Sheridan, M. A., & Lambert, H. K. (2014). Childhood adversity and neural development: Deprivation and threat as distinct dimensions of early experience. Neuroscience and Biobehavioral Reviews, 47, 578–591. https://doi.org/10.1016/j.neubiorev.2014.10.012

Merz, E. C., Wiltshire, C. A., & Noble, K. G. (2019). Socioeconomic Inequality and the Developing Brain: Spotlight on Language and Executive Function. Child Development Perspectives, 13(1), 15–20. https://doi.org/10.1111/cdep.12305

Miller-Cotto, D., Smith, L. V., Wang, A. H., & Ribner, A. D. (2021). Changing the conversation: A culturally responsive perspective on executive functions, minoritized children and their families. Infant and Child Development, April, 1–12. https://doi.org/10.1002/icd.2286

Orfield, G., & Lee, C. (2005). Why segregation matters: Poverty and educational inequality. The Civil Rights Project.

Owens, M. M., Yuan, D., Hahn, S., Albaugh, M., Allgaier, N., Chaarani, B., … & Garavan, H. (2020). Investigation of Psychiatric and Neuropsychological Correlates of Default Mode Network and Dorsal Attention Network Anticorrelation in Children. Cerebral Cortex, 30(12), 6083–6096.

Pines, A. R., Larsen, B., Cui, Z., Sydnor, V. J., Bertolero, M. A., Adebimpe, A., Alexander-Bloch, A. F., Davatzikos, C., Fair, D. A., Gur, R. C., Gur, R. E., Li, H., Milham, M. P., Moore, T. M., Murtha, K., Parkes, L., Thompson-Schill, S. L., Shanmugan, S., Shinohara, R. T., … Satterthwaite, T. D. (2021). Dissociable Multi-scale Patterns of Development in Personalized Brain Networks. BioRxiv, 2021.07.07.451458. https://doi.org/10.1101/2021.07.07.451458

Power, J. D., Mitra, A., Laumann, T. O., Snyder, A. Z., Schlaggar, B. L., & Petersen, S. E. (2014). Methods to detect, characterize, and remove motion artifact in resting state fMRI. NeuroImage. https://doi.org/10.1016/j.neuroimage.2013.08.048

Power, J. D., Schlaggar, B. L., & Petersen, S. E. (2014). Primer Studying Brain Organization via Spontaneous fMRI Signal. Neuron, 84(4), 681–696. https://doi.org/10.1016/j.neuron.2014.09.007

Power, J. D., Schlaggar, B. L., & Petersen, S. E. (2015). Recent progress and outstanding issues in motion correction in resting state fMRI. In NeuroImage. https://doi.org/10.1016/j.neuroimage.2014.10.044

R Core Team. (2017). R: A language and environment for statistical computing. R Foundation for Statistical Computing. https://www.r-project.org/

Raichle, M. E., MacLeod, A. M., Snyder, A. Z., Powers, W. J., Gusnard, D. A., & Shulman, G. L. (2001). A default mode of brain function. Proceedings of the National Academy of Sciences of the United States of America, 98(2), 676–682. https://doi.org/10.1073/pnas.98.2.676

Rakesh, D., Seguin, C., Zalesky, A., Cropley, V., & Whittle, S. (2021). Associations between neighborhood disadvantage, resting-state functional connectivity, and behavior in the Adolescent Brain Cognitive Development (ABCD) Study ® : Moderating role of positive family and school environments. Biological Psychiatry: Cognitive Neuroscience and Neuroimaging. https://doi.org/https://doi.org/10.1016/j.bpsc.2021.03.008

Rakesh, D., Zalesky, A., & Whittle, S. (2021). Similar but distinct – Effects of different socioeconomic indicators on resting state functional connectivity: findings from the Adolescent Brain Cognitive Development (ABCD) Study®. Developmental Cognitive Neuroscience, 51, 101005. https://doi.org/10.1016/j.dcn.2021.101005

Reardon, S. F., & Owens, A. (2014). 60 Years After Brown: Trends and Consequences of School Segregation. Annual Review of Sociology, 40(1), 199–218. https://doi.org/10.1146/annurev-soc-071913-043152

Satterthwaite, T. D., Wolf, D. H., Ruparel, K., Erus, G., Elliott, M. A., Eickhoff, S. B., Gennatas, E. D., Jackson, C., Prabhakaran, K., & Smith, A. (2013). Heterogeneous impact of motion on fundamental patterns of developmental changes in functional connectivity during youth. Neuroimage, 83, 45–57.

Scott, J. C., Pinderhughes, E. E., & Johnson, S. K. (2020). How does racial context matter?: family preparation-for-bias messages and racial coping reported by black youth. Child Development, 91(5), 1471–1490.

Seitzman, B. A., Gratton, C., Laumann, T. O., Gordon, E. M., Adeyemo, B., Dworetsky, A., Kraus, B. T., Gilmore, A. W., Berg, J. J., Ortega, M., Nguyen, A., Greene, D. J., McDermott, K. B., Nelson, S. M., Lessov-Schlaggar, C. N., Schlaggar, B. L., Dosenbach, N. U. F., & Petersen, S. E. (2019). Trait-like variants in human functional brain networks. Proceedings of the National Academy of Sciences of the United States of America, 116(45), 22851–22861. https://doi.org/10.1073/pnas.1902932116

Seli, P., Carriere, J. S. A., & Smilek, D. (2015). Not all mind wandering is created equal: dissociating deliberate from spontaneous mind wandering. Psychological Research, 79(5), 750–758. https://doi.org/10.1007/s00426-014-0617-x

Shedd, C. (2015). Unequal city: Race, schools, and perceptions of injustice. Russell Sage Foundation.

Sheridan, M. A., Sarsour, K., Jutte, D., D’Esposito, M., & Boyce, W. T. (2012). The impact of social disparity on prefrontal function in childhood. PLoS ONE, 7(4). https://doi.org/10.1371/journal.pone.0035744

Sherman, L. E., Rudie, J. D., Pfeifer, J. H., Masten, C. L., McNealy, K., & Dapretto, M. (2014). Development of the Default Mode and Central Executive Networks across early adolescence: A longitudinal study. Developmental Cognitive Neuroscience. https://doi.org/10.1016/j.dcn.2014.08.002

Spreng, R. N. (2012). The fallacy of a “task-negative” network. Frontiers in Psychology, 3(MAY), 1–5. https://doi.org/10.3389/fpsyg.2012.00145

Spreng, R. N., & Turner, G. R. (2019). The Shifting Architecture of Cognition and Brain Function in Older Adulthood. Perspectives on Psychological Science, 14(4), 523–542. https://doi.org/10.1177/1745691619827511

Sridharan, D., Levitin, D. J., & Menon, V. (2008). A critical role for the right fronto-insular cortex in switching between central-executive and default-mode networks. Proceedings of the National Academy of Sciences of the United States of America, 105(34), 12569–12574. https://doi.org/10.1073/pnas.0800005105

St Clair-Thompson, H. L., & Gathercole, S. E. (2006). Executive functions and achievements in school: Shifting, updating, inhibition, and working memory. Quarterly Journal of Experimental Psychology, 59(4), 745–759.

Tooley, U. A., Bassett, D. S., & Mackey, A. P. (2021). Environmental influences on the pace of brain development. Nature Reviews Neuroscience, 22(6), 372–384. https://doi.org/10.1038/s41583-021-00457-5

Uddin, L. Q., Supekar, K. S., Ryali, S., & Menon, V. (2011). Dynamic reconfiguration of structural and functional connectivity across core neurocognitive brain networks with development. Journal of Neuroscience, 31(50), 18578–18589. https://doi.org/10.1523/JNEUROSCI.4465-11.2011

Vincent, J. L., Kahn, I., Snyder, A. Z., Raichle, M. E., & Buckner, R. L. (2008). Evidence for a frontoparietal control system revealed by intrinsic functional connectivity. Journal of Neurophysiology, 100(6), 3328–3342. https://doi.org/10.1152/jn.90355.2008

Wechsler, D. (2014). Wechsler intelligence scale for children, 5th edition. Pearson.

Weissman, D. G., Hatzenbuehler, M. L., Cikara, M., Barch, D., & McLaughlin, K. A. (2021). Antipoverty programs mitigate socioeconomic disparities in brain structure and psychopathology among US youths. Psyarxiv. https://doi.org/10.31234/osf.io/8nhej

Weissman, D. H., Roberts, K. C., Visscher, K. M., & Woldorff, M. G. (2006). The neural bases of momentary lapses in attention. Nature Neuroscience, 9(7), 971–978. https://doi.org/10.1038/nn1727

Whitfield-Gabrieli, S., Wendelken, C., Nieto-Castañón, A., Bailey, S. K., Anteraper, S. A., Lee, Y. J., Chai, X. Q., Hirshfeld-Becker, D. R., Biederman, J., Cutting, L. E., & Bunge, S. A. (2020). Association of Intrinsic Brain Architecture with Changes in Attentional and Mood Symptoms during Development. JAMA Psychiatry, 77(4), 378–386. https://doi.org/10.1001/jamapsychiatry.2019.4208

Willoughby, M. T., Wylie, A. C., & Little, M. H. (2019). Testing longitudinal associations between executive function and academic achievement. Developmental Psychology, 55(4), 767.

Zucman, G. (2019). Global Wealth Inequality. Annual Review of Economics, 11, 109–138. https://doi.org/10.1146/annurev-economics-080218-025852

