## Supplementary materials for "Exploring neural correlates of behavioral and academic resilience among children in poverty"

### Supplemental materials for "Exploring neural correlates of behavioral and academic resilience among children in poverty"

### Analytic plan and results

Supplementary Table 1. A summary of our pre-registered aims, hypotheses, statistical test, and outcome for each of the tests we pre-registered. Tests are in R notation: The variable before ~ signals the DV, and those after signal IVs. X*Y indicates the inclusion of an interaction and main effect for variables X and Y, while the + indicates only a main effect. Finally the (1|Z) notation denotes the inclusion of random intercepts for Z.

| Aim 1: Network connectivity change over adolescence | | | |
|---|---|---|---|
| | **Hypothesis** | **Test** | **Outcome** |
| Primary | Decline in LFPN-DMN connectivity between T0 and T2; no interaction by poverty status | LFPN-DMN connectivity ~ timepoint [T0, T2] * poverty status [above, below] + motion + sex + (1\|id) + (1\|family) + (1\|site) | **Main effect**: not significant<br>**Interaction**: not significant |
| Secondary | Decline in CON-DMN connectivity between T0 and T2; possible weaker effect for children in poverty | CON-DMN connectivity ~ timepoint [T0, T2] * poverty status [above, below] + motion + sex + (1\|id) + (1\|family) + (1\|site) | **Main effect**: decline in CON-DMN connectivity<br>**Interaction**: weaker effect for children in poverty |
| Secondary | Decline in CON-LFPN connectivity between T0 and T2; possible weaker effect for children in poverty | CON-LFPN connectivity ~ timepoint [T0, T2] * poverty status [above, below] + motion + sex + (1\|id) + (1\|family) + (1\|site) | **Main effect**: decline in CON-LFPN connectivity<br>**Interaction**: not significant |

| Aim 2: Links between test performance and grades in school | | | |
|---|---|---|---|
| | **Hypothesis** | **Test** | **Outcome** |
| Primary | Positive association between children's cognitive test performance and their grades in school concurrently at T0; weaker for children in poverty | Grades at T0 ~ test performance at T0 * poverty status [above, below] + sex + (1\|family) + (1\|site) | **Main effect**: positive association<br>**Interaction**: weaker effect for children in poverty |
| Primary | Positive association between children's cognitive test at T0 performance and their grades in school at T2; weaker for children in poverty | Grades at T2 ~ test performance at T0 * poverty status [above, below] + grades at T0 + sex + (1\|family) + (1\|site) | **Main effect**: positive association<br>**Interaction**: not significant |

| Aim 3: Links between network connectivity, grades in school, and attention problems | | | |
|---|---|---|---|
| | **Hypothesis** | **Test** | **Outcome** |
| Primary | Differential association between LFPN-DMN connectivity and grades at T0 for children above vs below poverty (above: higher LFPN-DMN~worse grades; below: higher LFPN-DMN~better grades) | Grades at T0 ~ LFPN-DMN connectivity at T0 * poverty status [above, below] + motion + sex + (1\|family) + (1\|site) | **Main effect**: higher LFPN-DMN associated with worse grades<br>**Interaction**: differential association for children above and below poverty (above: higher LFPN-DMN~worse grades; below: higher LFPN-DMN~better grades) |
| Primary | Longitudinal association between LFPN-DMN connectivity at T0 and grades at T2 (above: positive; below: negative) | Grades at T2 ~ LFPN-DMN connectivity at T0 * poverty status [above, below] + grades at T0 + motion + sex + (1\|family) + (1\|site) | **Main effect**: not significant<br>**Interaction**: not significant |
| Primary | Positive association between LFPN-DMN connectivity at T0 and attention problems at T2; possible for the reverse for children in poverty | Attention problems at T2 ~ LFPN-DMN connectivity at T0 * poverty status [above, below] + attention problems at T0 + motion + sex + (1\|family) + (1\|site) | **Main effect**: not significant<br>**Interaction**: not significant |
| Secondary | Negative association between CON-LFPN connectivity and academic performance at T0; no interaction by poverty status | Grades at T0 ~ CON-LFPN connectivity at T0 * poverty status [above, below] + motion + sex + (1\|family) + (1\|site) | **Main effect**: higher CON-LFPN associated with worse grades<br>**Interaction**: not significant |
| Secondary | Positive association for both CON-LFPN and CON-DMN connectivity and attention problems at T0; stronger for children below poverty | Grades at T0 ~ LFPN-DMN connectivity at T0 * poverty status [above, below] + CON-DMN connectivity at T0 * poverty status [above, below] + CON-LFPN connectivity at T0 * poverty status [above, below] + motion + sex + (1\|family) + (1\|site) | **Main effect**:<br>higher CON-DMN connectivity associated with more attention problems<br>CON-LFPN not significant<br>**Interaction**:<br>stronger CON-DMN association for children below poverty<br>CON-LFPN not significant |

### Secondary aims

**Secondary Aim 1**. Parallel to Primary Aim 1, we also sought to characterize resting state coupling changes for CON-DMN and CON-LFPN over early adolescence. Based on prior findings from the literature (e.g., Grayson & Fair, 2017), these networks are expected to become increasingly decoupled during development. If this is a general phenomenon (secondary H1a), we expected to find decreased connectivity for these networks for both children above and below poverty. However, because we found previously that children below poverty with higher test scores tended to have low DMN-CON connectivity at T0, it is possible that some children in poverty may not show a reduction in coupling with age, leading to a weaker effect for the group as a whole (secondary H1b).

**Secondary Aim 2**. Parallel to Primary Aim 3, we also sought to test whether CON-LFPN and CON-DMN were associated with grades and attention problems.

*CON-LFPN associations with academic performance*: We hypothesized that stronger CON-LFPN connectivity would be associated with worse academic performance in both children above and below poverty (secondary H2). We planned to test this hypothesis concurrently.

*Interplay between LFPN-DMN, CON-LFPN, and CON-DMN and attention problems*: Though immature at age 9, CON is theorized to serve as an intermediary the DMN and LFPN, enabling switching attention between internally and externally guided mental states. Thus, we planned to test whether these patterns of CON connectivity were differentially associated with attention problems for children above and below poverty. We predicted that weaker CON-LFPN and CON-DMN connectivity would be associated with fewer attention problems for children below (but not above) poverty (secondary H3). We planned to test this hypothesis concurrently.

### Sample exclusion flowchart

The charts below illustrates how many children were excluded sequentially with each additional exclusion criteria (i.e., after excluding children with missing rs-fMRI data, an additional 857 children had missing income data), at T0 (immediately below), and at T2 (next page).

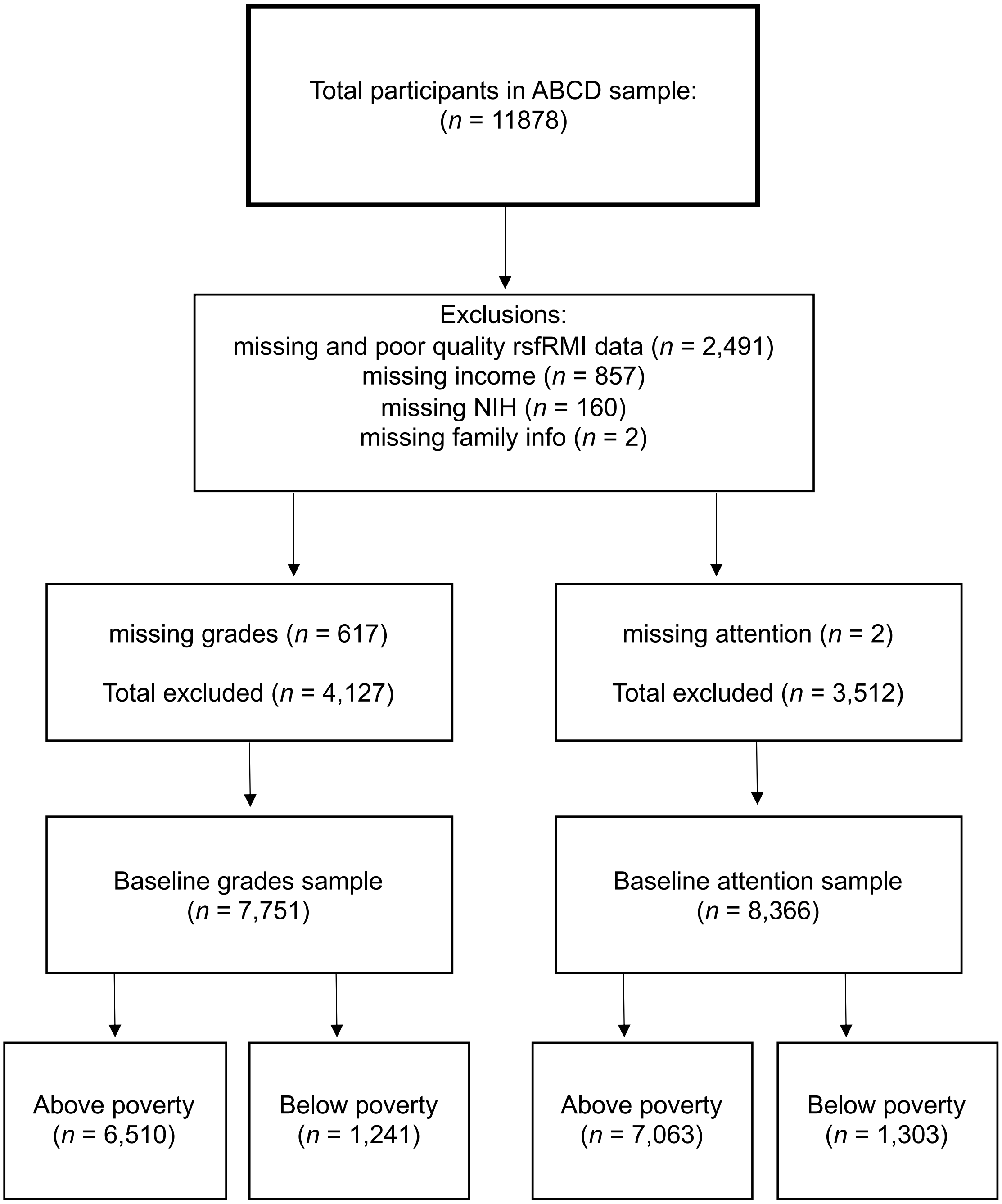

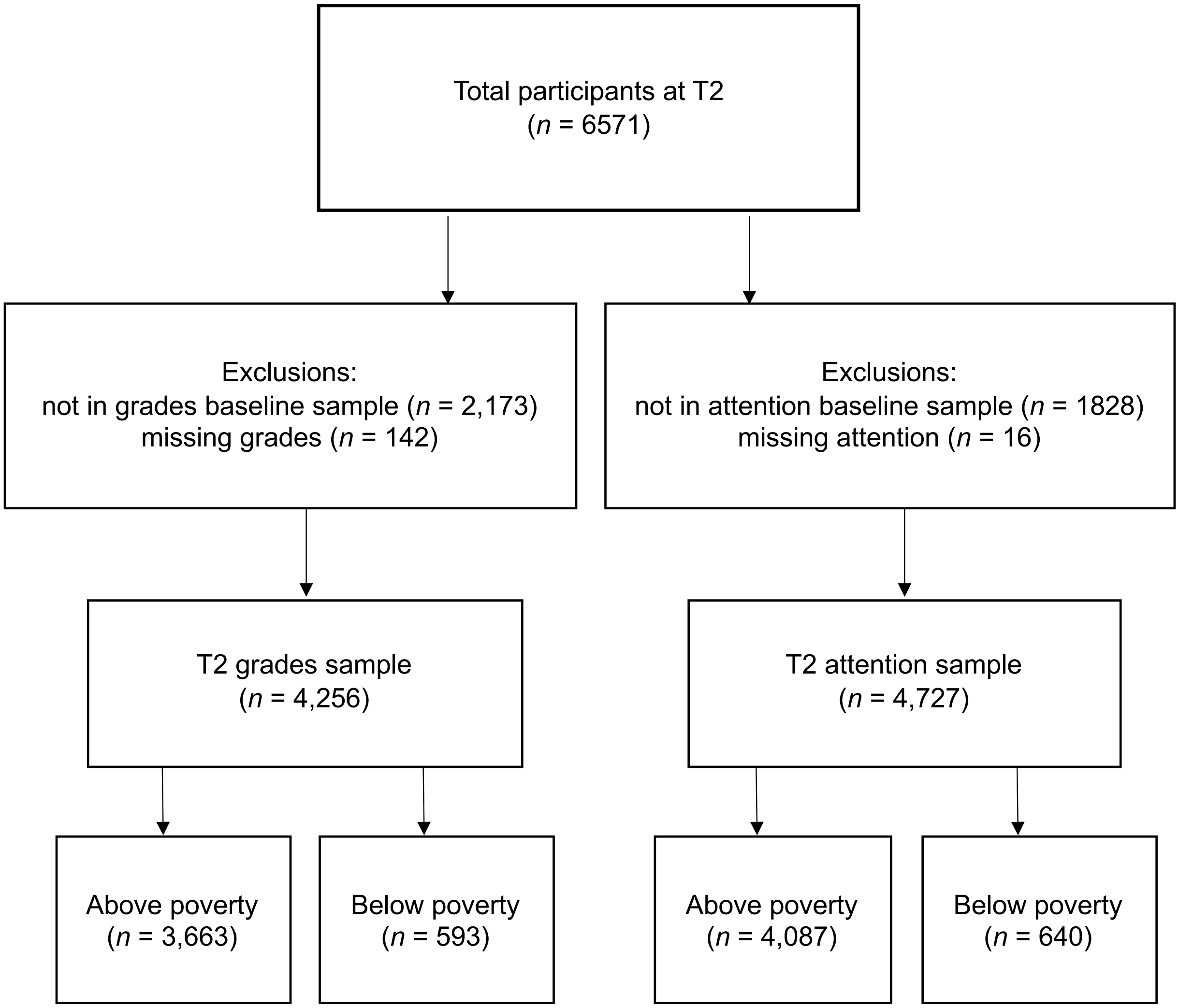
Total participants at T2
(n = 6571)
Exclusions:
not in grades baseline sample (n = 2,173)
missing grades (n = 142)
Exclusions:
not in attention baseline sample (n = 1828)
missing attention (n = 16)
T2 grades sample
(n = 4,256)
T2 attention sample
(n = 4,727)
Above poverty
(n = 3,663)
Below poverty
(n = 593)
Above poverty
(n = 4,087)
Below poverty
(n = 640)

### Differences between children excluded and included in sample

The children who were excluded differed meaningfully from our final sample. In particular, they were more likely to be in poverty, be younger, be male, have more attention problems, have lower grades, and have lower cognitive test scores, as displayed in Supplementary Table 2 below.

| | Included | Excluded | p test |
|---|---|---|---|
| *n* | 7751 | 4109 | |
| Age (mean (SD)) | 9.95 (0.63) | 9.85 (0.61) | <0.001 |
| Sex = M (%) | 3850 (49.7) | 2335 (56.8) | <0.001 |
| Income = below poverty (%) | 1241 (16.0) | 584 (19.7) | <0.001 |
| Grades (mean (SD)) | 1.61 (0.75) | 1.87 (0.86) | <0.001 |
| Attention (mean (SD)) | 53.52 (5.87) | 54.63 (6.63) | <0.001 |
| NIH test scores (mean (SD)) | 92.76 (10.20) | 89.13 (11.12) | <0.001 |

Supplementary Table 2. Differences between excluded and included participants in variables of interest, using the sample of children with usable grades data. Mean (SD) for variables for the two groups displayed; p-value represents the significance of a two-sided t-test.

Focusing specifically on whether children had usable rs-fMRI data revealed the same patterns, suggesting this may the limiting factor in maintaining a representative sample (Supplementary Table 3).

| | Included | Excluded | p test |
|---|---|---|---|
| *n* | 9387 | 2390 | |
| Age (mean (SD)) | 9.94 (0.63) | 9.81 (0.60) | <0.001 |
| Sex = M (%) | 4698 (50.0) | 1452 (60.8) | <0.001 |
| Income = below poverty (%) | 1341 (15.7) | 464 (21.9) | <0.001 |
| Grades (mean (SD)) | 1.63 (0.77) | 1.88 (0.87) | <0.001 |
| Attention (mean (SD)) | 53.62 (5.95) | 54.97 (6.83) | <0.001 |
| NIH test scores (mean (SD)) | 92.36 (10.31) | 88.61 (11.27) | <0.001 |

Supplementary Table 3. Differences between participants in the entire sample who did and did not meet criteria for usable rs-fMRI data. Mean (SD) for variables for the two groups displayed; p-value represents the significance of a two-sided t-test.

While there were proportionally more children in poverty excluded, it is important to note that these differences in age, sex, attention problems, grades, and cognitive test scores are present between excluded and included children both above and below poverty (Supplementary Table 4).

| | Above poverty | | Below poverty | | |
|---|---|---|---|---|---|
| | included | excluded | included | excluded | p test |
| *n* | 6510 | 2379 | 1241 | 584 | |
| Age (mean (SD)) | 9.96 (0.63) | 9.82 (0.61) | 9.90 (0.62) | 9.83 (0.60) | <0.001 |
| Sex = M (%) | 3233 (49.7) | 1396 (58.7) | 617 ( 49.7) | 319 ( 54.6) | <0.001 |
| Grades (mean (SD)) | 1.53 (0.71) | 1.74 (0.81) | 2.02 (0.87) | 2.21 (0.95) | <0.001 |
| Attention (mean(SD)) | 53.29 (5.55) | 54.51 (6.37) | 54.71 (7.17) | 55.78 (7.49) | <0.001 |
| NIH test scores (mean (SD)) | 93.80 (9.75) | 90.92 (10.63) | 87.32 (10.75) | 84.41 (11.27) | <0.001 |

Supplementary Table 4. Differences between excluded and included participants in variables of interest, using the sample of children with usable grades data, divided by whether children are above or below poverty. Mean (SD) for variables for the four groups displayed; p-value represents the significance of a two-sided t-test between the excluded and included samples.

Thus, regardless of whether or not children are in poverty, demands associated with the study—in particular, producing usable rs-fMRI data—selected for children who were older, female, and higher-performing.

### Characterizing behavioral measures

Most children in our sample were reported by their parents as receiving A's and B's, although the broader range from A-F was represented. On average, children below poverty received lower grades ($M$ = 2.02, SD = 0.87) than children above poverty ($M$ = 1.53, SD = 0.71). This difference in grades between children living below and above poverty was significant ($t$(1569.1) = -18.51, $p < 0.001$). In terms of attention, most children in our sample scored below the clinical range on the CBCL attention subscale at T0. On average, children in poverty had more attention problems ($M$ = 54.80, SD = 7.30) than children above poverty ($M$ = 53.36, SD = 5.58). This difference in attention scores between children living below and above poverty was also significant ($t$(1594.7) = -6.80, $p < 0.001$).

All three behavioral measures—NIH cognitive test scores, grades, and attention—were significantly correlated with one another, although there was a range of r-values between concurrently tested variables (Supplementary Figure 1). Notably, attention was relatively weakly correlated with NIH test scores ($r$ = .14 at T0, the timepoint at which NIH test scores were available) but moderately concurrently related to grades ($r$ = .36-.38 across three timepoints). Grades and NIH test scores were also moderately correlated ($r$ = .33 at baseline). Thus, although these three variables were interrelated, they were by no means redundant with one another. Additionally, we found that attention was fairly stable over the three timepoints ($r$ = .68 and .60 for T0 vs. the 1-year and 2-year follow-ups, respectively). So, too, were parent-reported grades ($r$ = .74 and .68).

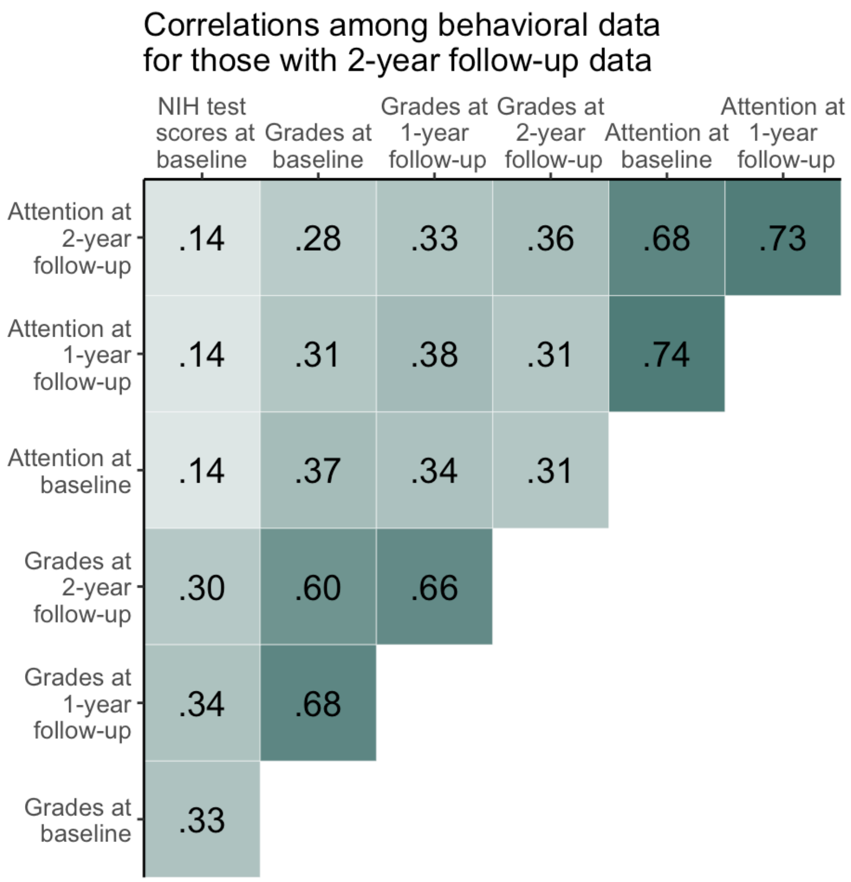

**Supplementary Figure 1.** R-values for correlations among behavioral measures used in the current study, for children with data at all three timepoints. For the purposes of correlations, grades are treated as a continuous rather than categorical variable; all variables have also been recoded so that higher scores indicate better functioning (higher grades, fewer attention problems). All correlations are significant to the $p <$ .0001 level.

### Longitudinal changes in network development

We also examined changes in functional connectivity for CON-DMN and CON-LFPN, as secondary pre-registered analyses. Based on prior studies, we expected that these networks would become increasingly decoupled during development, for both children above and below poverty. We further predicted this effect may be attenuated for children below poverty. Results are displayed in Supplementary Figure 2 B and C.

**CON-DMN connectivity over adolescence**. On average, CON-DMN connectivity decreased significantly over ages 9-13, confirming our prediction, $B$ = -0.01, SD = 0.001, $\chi^2$ (2) = 61.55, $p$ < .001. Notably, this relation interacted significantly as a function of poverty status, interaction: $B$ = -0.001, SD = 0.002, $\chi^2$ (1) = 6.61, $p$ = .010 (Supplementary Figure 2). While children below poverty did not show a significant decrease in connectivity, this change was significant for children above poverty (below poverty: $B$ = -0.004, SD = 0.003, $\chi^2$ (1) = 1.52, $p$ = .217; above poverty: $B$ = -0.009, SD = 0.001, $\chi^2$ (1) = 56.47, $p$ < .001).

**CON-LFPN connectivity over adolescence**. Similarly, CON-LFPN connectivity decreased significantly across the two years, on average, $B$ = -0.005, SD = 0.001, $\chi^2$ (2) = 19.28, $p$ < .001. This main effect was qualified by a non-significant interaction as a function of poverty status, interaction: $B$ = 0.005, SD = 0.003, $\chi^2$ (1) = 3.55, $p$ = .060 (Supplementary Figure 2).

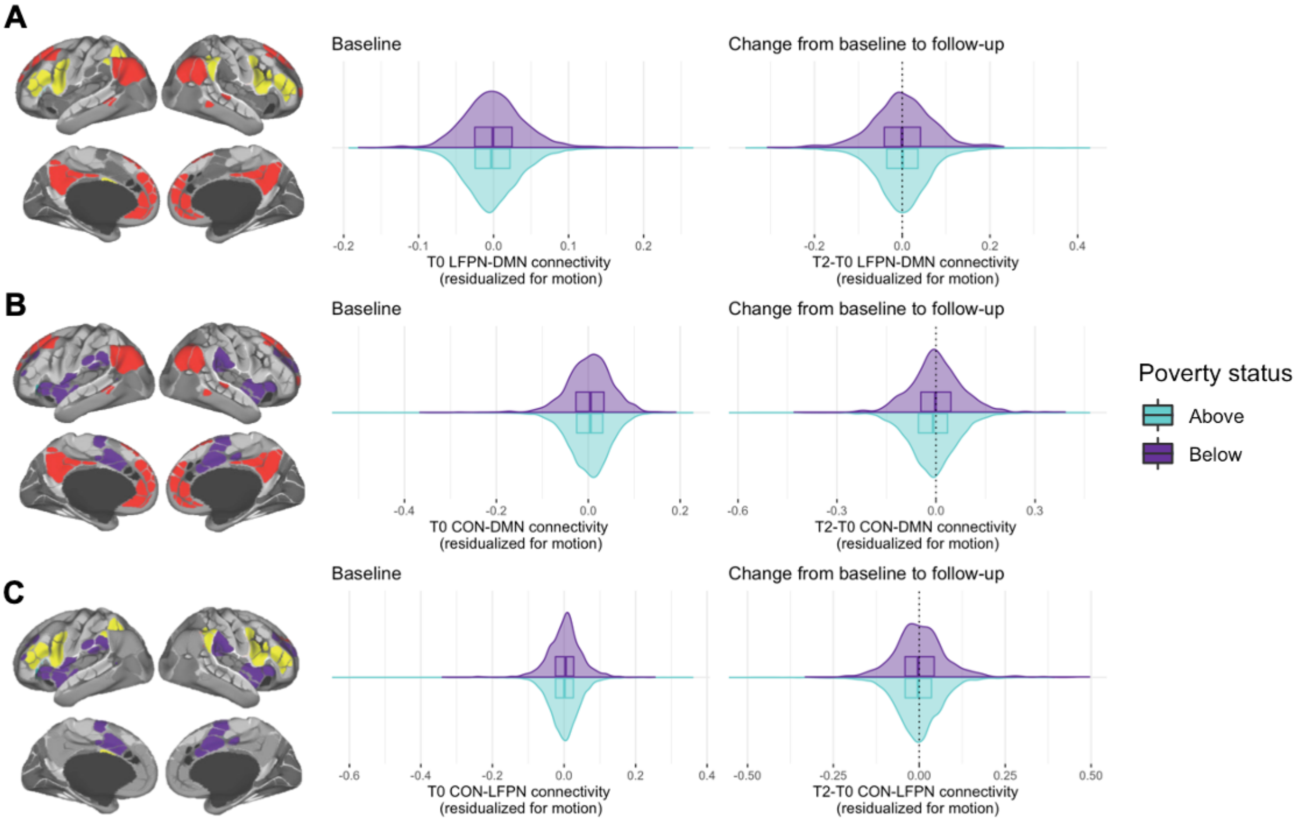

Supplementary Figure 2. Connectivity for LFPN-DMN (**A**), CON-DMN (**B**), and CON-LFPN (**C**). Distribution of values is displayed as both frequency plots and box plots; the center panel displays connectivity values at T0 after residualizing for motion, while the right panel displays the difference in residualized connectivity between T0 and T2. In right panel, dotted line at zero indicates no change; negative values indicate a decrease between T0 and T2. Lighter teal color (inverted frequency plots) indicates children above poverty, while purple (top frequency plots) indicates children below poverty.

### Longitudinal network development: interactions with test performance

***Developmental trajectories of networks as a function of poverty status and test performance***

As exploratory analyses, we investigated whether trajectories of network connectivity differ as a function of children's cognitive test scores and their poverty status. To this end, we conducted three separate linear mixed effects models associating (1) LFPN-DMN connectivity at T2, (2) CON-DMN connectivity at T2, and (3) CON-LFPN connectivity at T2, respectively, with a three-way interaction between connectivity at T0, poverty status, and T0 cognitive test performance.

***LFPN-DMN trajectories and test scores.*** LFPN-DMN connectivity at T0 was predictive of connectivity two years later, $B$ = 0.29, SD = 0.164, $\chi^2$ (4) = 334.78, $p$ < .001. There were no significant interactions.

***CON-DMN trajectories and test scores.*** CON-DMN connectivity at T0 was predictive of CON-DMN connectivity two years later, $B$ = 0.08, SD = 0.156, $\chi^2$ (4) = 306.13, $p$ < .001. There were no significant interactions.

***CON-LFPN trajectories and test scores.*** CON-LFPN connectivity at T0 was predictive of CON-LFPN connectivity two years later, $B$ = 0.01, SD = 0.158, $\chi^2$ (4) = 269.55, $p$ < .001. There was also a significant interaction of CON-LFPN network connectivity at T0 by poverty status, $B$ = 0.04, SD = 0.355, $\chi^2$ (4) = 24.86, $p$ < .001. The relation was significant for both children below and above poverty, though it was in opposite directions (below poverty: $B$ = -0.035, SD = 0.319, $\chi^2$ (2) = 8.87, $p$ = 0.012; above poverty: $B$ = 0.015, SD = 0.158, $\chi^2$ (2) = 262.85, $p$ < .001). Thus, children below poverty who started out with higher CON-LFPN connectivity at T0 showed lower CON-LFPN connectivity at T2, while children above poverty who had higher CON-LFPN connectivity at T0 also showed higher CON-LFPN connectivity at T2.

Thus, across all networks, children's test scores at T0 were not associated with the rate of change in their connectivity metrics over the next two years, regardless of children's poverty status.

### Relations between academic performance and CON-LFPN connectivity

Because CON has been posited to play a role in alerting LFPN to external challenges, we conducted a preregistered secondary analysis testing whether CON-LFPN connectivity was concurrently related to children's grades. Based on our prior work in this sample examining cognitive test performance, we predicted that stronger CON-LFPN connectivity would be associated with worse academic performance in both children above and below poverty.

**Secondary analyses**. Indeed, we found that on average, higher CON-LFPN connectivity was related to worse grades concurrently at T0, *B* = 0.71, SD = 0.48, $\chi^2$ (2) = 6.70, *p* = .035. However, this relation did not differ as a function of poverty status, interaction: *B* = -3.11, SD = 1.11, $\chi^2$ (1) = 1.51, *p* = .219; therefore, we did not follow up with exploratory longitudinal analyses. This finding suggests that CON-LFPN segregation may generally be beneficial for academic performance.

### Contribution of CON connectivity to attention problems

CON has been theorized to serve as an intermediary between DMN and LFPN, enabling switching attention between internally and externally guided mental status. Thus, as secondary analyses, we tested whether patterns of CON connectivity at baseline were differentially associated with attention problems for children above and below poverty, after accounting for LFPN-DMN connectivity and its interaction with poverty status.

The output of the model is displayed in Supplementary Table 5. In this model including all three network pairings, higher CON-DMN connectivity was related to worse attention on average ($B$ = 3.48, SD = 1.15, $\chi^2$ (1) = 20.86, $p$ < .001). In addition, both CON-DMN and LFPN-DMN connectivity differed significantly as a function of poverty status (CON-DMN by poverty interaction: $B$ = 7.51, SD = 2.68, $\chi^2$ (1) = 7.85, $p$ = .005; LFPN-DMN by poverty interaction: $B$ = -8.26, SD = 3.07, $\chi^2$ (1) = 7.24, $p$ = .007). Thus, CON-DMN connectivity and its association with poverty status was associated with attention problems over and above the interaction of LFPN-DMN connectivity with poverty status, which also continued to add significant variance; CON-LFPN connectivity did not contribute significant variance in this model.

We next performed exploratory follow-up analyses breaking down these interactions. As in our previous analyses focused solely on LFPN-DMN, this model with all three networks showed a positive relation between LFPN-DMN connectivity and attention problems for children above poverty, but a negative, non-significant relation for children below poverty (above poverty: $B$ = 3.53, SD = 1.25, $\chi^2$ (1) = 7.91, $p$ = 0.005, below poverty: $B$ = -4.02, SD = 3.47, $\chi^2$ (1) = 1.36, $p$ = 0.243). On the other hand, higher CON-DMN connectivity was associated with more attention problems for both children above and below poverty—and the association was in fact stronger for children below poverty (above poverty: $B$ = 3.43, SD = 1.11, $\chi^2$ (1) = 9.58, $p$ = 0.002, below poverty: $B$ = 10.14, SD = 3.03, $\chi^2$ (1) = 11.14, $p$ = 0.001). Indeed, the strongest concurrent predictor of attention problems among children below poverty was high CON-DMN connectivity.

Additional exploratory follow-up analyses focusing only on CON-DMN showed that baseline CON-DMN was not predictive of future attention problems when controlling for initial attention scores. However, when not controlling for initial attention scores, there was a significant interaction between baseline CON-DMN connectivity and poverty status in predicting T2 attention (interaction at T2: $B$ = 11.08, SE = 3,12, $\chi^2$ (1) = 12.58, $p$ < .001). Mirroring our results at T0, follow-up analyses showed that the longitudinal relation was significant for children below poverty, but not above poverty (below poverty: $B$ = 12.17, SD = 3.31, $\chi^2$ (1) = 13.41, p < 0.001; above poverty: B = 0.81, SD = 1.26, $\chi^2$ (1) = 0.41, p = 0.522). Thus, high CON-DMN connectivity at baseline was associated with attention problems for children below poverty at two timepoints.

| Attention | *B* | SD | $\chi^2$ | *p* |
|---|---|---|---|---|
| **LFPN-DMN** | 3.47 | 1.31 | 2.82 | 0.093 |
| **CON-DMN** | 3.48 | 1.15 | 20.86 | < 0.001 *** |
| **CON-LFPN** | -0.48 | 1.33 | 0.051 | 0.821 |
| **Poverty level** | 2.47 | 0.38 | 40.27 | < 0.001 *** |
| **Motion** | 1.71 | 0.39 | 19.40 | < 0.001 *** |
| **Sex(M)** | 0.36 | 0.13 | 7.62 | 0.006 ** |
| **LFPN-DMN*poverty level** | -8.26 | 3.07 | 7.24 | 0.007 ** |
| **CON-DMN*poverty level** | 7.51 | 2.68 | 7.85 | 0.005 ** |
| **CON-LFPN*poverty level** | 4.08 | 3.07 | 1.77 | 0.184 |

Supplementary Table 5**.** Results of linear mixed effects model associating attention problems at T0 with an interaction between each of the three brain networks of interest (LFPN-DMN, CON-DMN, and CON-LFPN), separately, and poverty status. Chi-squared and significance values from Type II anova, using the Anova package in *car* (Fox & Weisberg, 2019)

### Participant information and analyses at the T1 timepoint

The table below summarizes participant characteristics at T1. Next, we report supplementary longitudinal analyses with regard to grades and attention scores at T1.

| **Timepoint** | | **Attention data (*N* = 8366)** | | **Grades data (*N* = 7751)** | |
|---|---|---|---|---|---|
| | | Above poverty | Below poverty | Above poverty | Below poverty |
| **One-year follow-up (T1)** | *N* | 6780 | 1155 | 5984 | 1060 |
| | Sex | F: 3386<br>M: 3394 | F: 577<br>M: 578 | F: 3018<br>M: 2966 | F: 528<br>M: 532 |
| | Age | 9.1-12.4 | 9.1-12.4 | 9.7-12.4 | 9.7-12.4 |

Supplementary Table 6. Sample sizes, age, and parent-reported child sex for children above and below poverty at T1. Sample sizes differ slightly for those analyses focusing on grades and those focusing on attention, based on the number of children providing usable data.

**Associations between grades and cognitive test scores**. Higher cognitive test scores were related to higher grades in school one year later, controlling for grades at T0, B = -0.05, SD = 0.004, $\chi$2 (2) = 174.99, p < .001. However, in contrast to our prediction, this relation did not differ significantly as a function of poverty status, interaction: B = 0.01, SD = 0.01, $\chi$2 (1) = 1.07, p = .301.

**Associations between grades and LFPN-DMN connectivity**. Higher LFPN-DMN connectivity was related to worse grades one year later, controlling for grades at T0, *B* = 1.78, SD = 0.61, $\chi^2$ (2) = 13.31, *p* = .001. However, this relation differed significantly as a function of poverty status, interaction: *B* = -4.35, SD = 1.35, $\chi^2$ (1) = 10.58, *p* = .001. Specifically, higher LFPN-DMN connectivity appeared to be related to worse grades for children above poverty (clmm did not converge, parameters from lmer were *B* = 0.34, SD = 0.12), but directionally related to better grades for children below poverty: *B* = -1.90, SD = 1.05, $\chi^2$ (1) = 3.33, *p* = .068.

**Associations between attention and LFPN-DMN connectivity.** Contrary to our prediction, we found no significant relation between LFPN-DMN and attention at T1 when controlling for attention at T0, *B* = -0.89, SD = 0.84, $\chi^2$ (2) = 1.49, *p* = .474. Further, this relation did not differ significantly as a function of poverty status at either timepoint, interaction: *B* = 1.98, SD = 2.02, $\chi^2$ (1) = 0.97, *p* = .326.

### Substituting matrix reasoning for the NIH composite

In our previous study, Matrix Reasoning had shown the strongest group interaction in the link between LFPN-DMN connectivity and cognition. Therefore, we performed each analysis reported that tested associations with NIH cognitive test performance, substituting NIH composite with Matrix Reasoning.

Children's performance was measured on the Matrix Reasoning Task from the Wechsler Intelligence Test for Children-V (WISC-V), a measure of abstract reasoning (Wechsler, 2014). We used the total score for each child, at T0. Matrix Reasoning is a widely used test of higher-level cognition that was not included in the NIH composite score.

***Relations between academic performance and matrix reasoning.*** On average, higher matrix reasoning was related to higher grades concurrently, $B$ = -0.24, SD = 0.01, $\chi^2$ (2) = 614.95, $p$ < .001, though this relation differed as a function of poverty status, $B$ = 0.06, SD = 0.02, $\chi^2$ (1) = 5.71, $p$ = .017. For both children above and below poverty, higher reasoning scores were related to higher grades, though the relation was stronger for children above poverty (above poverty: $B$ = -0.25, SD = 0.012, $\chi^2$ (1) = 539.04, $p$ = .001; below poverty: $B$ = -0.18, SD = 0.02, $\chi^2$ (1) = 74.77, $p$ = .001). These results mirror the primary results found between children's performance on cognitive tests and grades in school.

We also conducted these analyses longitudinally. The same pattern was found at T1, such that higher matrix reasoning was related to higher grades, controlling for grades at T0, $B$ = -0.14, SD = 0.01, $\chi^2$ (2) = 121.46, $p$ < .001. However, this relation differed significantly as a function of poverty status, just as with the NIH toolbox composite, interaction: $B$ = 0.07, SD = 0.03, $\chi^2$ (1) = 6.73, $p$ = 0.009. At T2, higher matrix reasoning was again related to higher grades, controlling for grades at T0, $B$ = -0.14, SD = 0.02, $\chi^2$ (2) = 84.02, $p$ < .001, however, this relation did not differ as a function of poverty status, interaction: $B$ = 0.05, SD = 0.04, $\chi^2$ (1) = 2.07, $p$ = 0.150.

***Relations between attention problems and matrix reasoning.*** On average, children with higher matrix reasoning scores had fewer attention problems, $B$ = -0.04, SD = 0.01, $\chi^2$ (2) = 47.27, $p$ < .001. This relation did not differ significantly as a function of poverty status, $B$ = 0.01, SD = 0.01, $\chi^2$ (1) = 0.72, $p$ = 0.396.

***Interaction between changes in connectivity, poverty status, and matrix reasoning.***

***LFPN-DMN trajectories and matrix reasoning.*** LFPN-DMN connectivity at T0 was predictive of connectivity two years later, $B$ = 0.25, SD = 0.063, $\chi^2$ (4) = 331.12, $p$ < .001. There were no significant interactions with poverty or matrix reasoning.

***CON-DMN trajectories and matrix reasoning.*** CON-DMN connectivity at T0 was predictive of connectivity two years later, $B$ = 0.28, SD = 0.061, $\chi^2$ (4) = 301.08, $p$ < .001. There were no significant interactions with poverty or matrix reasoning.

***CON-LFPN trajectories and matrix reasoning.*** There were several significant main effects and interactions, including a significant three-way interaction between

CON-LFPN, poverty status, and matrix reasoning. Model parameters and significance are displayed in Supplementary Table 7.

| CON-LFPN T2 | *B* | SD | $\chi^2$ | *p* |
|---|---|---|---|---|
| **CON-LFPN T0** | 0.30 | 0.06 | 258.67 | < 0.001 *** |
| **Poverty level** | -0.01 | 0.01 | 3.64 | 0.057 |
| **Matrix reasoning** | -0.00 | 0.00 | 0.95 | 0.331 |
| **Motion T2** | 0.05 | 0.01 | 70.49 | < 0.001 *** |
| **Motion T0** | 0.01 | 0.01 | 4.23 | 0.040 * |
| **Sex(M)** | 0.01 | 0.00 | 17.55 | < 0.001 *** |
| **CON-LFPN T0 * poverty status** | -0.52 | 0.13 | 17.12 | < 0.001 *** |
| **CON-LFPN T0 * matrix reasoning** | -0.00 | 0.01 | 0.65 | 0.422 |
| **Poverty level T0 * matrix reasoning** | 0.00 | 0.00 | 0.11 | 0.743 |
| **CON-LFPN T0 * poverty status * matrix reasoning** | 0.04 | 0.01 | 8.05 | 0.005 ** |

Supplementary Table 7**.** Results of linear mixed effects model associating CON-LFPN T2 network connectivity with a three-way interaction between connectivity at T1, poverty status, and matrix reasoning scores. Chi-squared and significance values from Type II anova, using the Anova function in *car* (Fox & Weisberg, 2019).

### Deviations from pre-registration.

**Vigilance.** We had initially intended to look at vigilance as a potential mechanism of resilience for children below poverty, as specified in our preregistration. More specifically, we sought to explore whether vigilance could explain why higher LFPN-DMN connectivity is related to higher cognitive test scores for these children. However, the only measure of vigilance available in the ABCD dataset was in the KSADS-5 Diagnostic Interview at T0, with two questions about past and present hypervigilance (1 = yes, 0 = no). This measure did not fully capture our definition of vigilance and the number of children who endorsed the two items was very low; therefore, we do not report any analyses involving vigilance.

**Grades.** We also pre-registered that we would use the mean of parent-reported grades from the CBCL for academic performance. Instead, we used parent-reported grades in the ABCD Longitudinal Parent Diagnostic Interview for DSM-5 Background Items Full (KSAD) because there was no CBCL question on grades in the ABCD dataset that was available.

In addition, we preregistered an analysis plan using linear mixed effects models to test these relations. However, because grades are a categorical ordered variable, cumulative link mixed models are more appropriate. Thus, as noted in the main text, we report the latter analyses for all tests including grades as an outcome variable. Results are not meaningfully different when performing the pre-registered linear mixed effects models.
